# ^31^P NMR Spectroscopy Demonstrates Large Amounts of Phosphohistidine in Mammalian Cells

**DOI:** 10.1101/2020.12.03.409540

**Authors:** Mehul V. Makwana, Sandra van Meurs, Andrea M. Hounslow, Mike P. Williamson, Richard F. W. Jackson, Richmond Muimo

## Abstract

Protein phosphorylation plays a key role in many cellular processes but there is presently no accurate information or reliable procedure to determine the relative abundance of many phosphoamino acids in cells. At pH ≤ 8, phosphohistidine is unstable compared to the extensively studied phosphoserine, phosphothreonine and phosphotyrosine. This study reports the absolute quantitative analysis of histidine phosphorylation of proteins from a human bronchial epithelial cell (16HBE14o-) lysate using ^31^P NMR spectroscopic analysis. The method was designed to minimize loss of the phosphohistidine phosphoryl group. Phosphohistidine was determined on average to be approximately one third as abundant as phosphoserine and phosphothreonine combined (and thus roughly 20 times more abundant than phosphotyrosine). The amount of phosphohistidine, and phosphoserine/phosphothreonine per gram of protein from a cell lysate was determined to be 23 μmol/g and 68 μmol/g respectively. The amount of phosphohistidine, and phosphoserine/phosphothreonine per cell was determined to be 1.8 fmol/cell, and 5.8 fmol/cell respectively. After tryptic digest of proteins from the16HBE14o- cell lysate, the phosphohistidine signal was abolished and increasing phosphoserine/phosphothreonine signal was observed, which has implications for mass spectrometry investigations. The ^31^P NMR spectroscopic analysis not only highlights the abundance of phosphohistidine, which likely reflects its importance in mammalian cells, but also provides a way of measuring and comparing levels of phosphorylated amino acids in cells.

## Introduction

Phosphorylation is one of the most common post-translational modifications (PTMs)^1^ and is involved in a vast number of cellular processes in many life forms. There are thought to be nine phosphorylated amino acid residues in nature: phosphoserine (pSer), phosphothreonine (pThr), phosphotyrosine (pTyr), phosphohistidine (pHis),^2^ phospholysine (pLys),^3^ phosphoarginine (pArg),^3^ phosphoaspartic acid (pAsp),^4^ phosphoglutamic acid (pGlu),^4^ and phosphocysteine (pCys).^5^ From a molecular perspective, phosphorylation of an amino acid residue results in a change in charge (the phosphoryl group exists mostly as a dianion under physiological conditions^6^) and thus a change in the surface potential of a protein.^7^ At the cellular level this can be considered a large change, and mis-regulated phosphorylation can be detrimental to biological function.^8, 9^

The existence of phosphorylated amino acid residues diversifies the role of phosphorylation in biology as each phosphorylated amino acid residue has different chemical properties. For example, the oxygen-phosphorus linked amino acids (pSer, pThr and pTyr) are relatively acid stable compared to the acid labile nitrogen-linked phosphoramidate (pHis, pLys, pArg), carboxy oxygen-linked acyl phosphate (pAsp, pGlu), and sulfur-linked phosphorthiolate (pCys).^5, 10^ Routinely used amino acid analysis methods employ acidic conditions^10^ which can destroy acid labile phosphorylation. Therefore, it is not surprising that pSer, pThr, and pTyr currently dominate the “Proteome Wide PTM Statistics Curator”^1^ and “PhosphoSitePlus”^11^ phosphorylation statistics, suggesting that acid labile phosphorylated amino acids are not as abundant or important and have largely not been detected (and may therefore have been neglected).^1^ To date the majority of protein phosphorylation research (especially in mammalian cells) has been focused on pSer, pThr and pTyr.

There is clear evidence for histidine (His) phosphorylation in nature,^2^ and mass spectrometry (MS) studies by Hardman *et al*. on HeLa cells^12^ and Potel *et al*. on *E. coli*^13^ suggest pHis may be more widespread in both eukaryotes and prokaryotes than previously thought. Unlike other phosphorylated amino acid residues, there are two isomers of pHis, namely τ-pHis and π-pHis (Supplementary Fig. S1). Several reviews have addressed the role of pHis in cells and conventional wisdom considers pHis as a high energy intermediate in the transfer of the phosphoryl group in intermediate or central metabolism and cell signalling.^2, 14–16^ However, there may be other roles of pHis in biology that have not yet been discovered. Therefore, the question remains as to how abundant is pHis (and other phosphorylated amino acids) in cells?

Determining the abundance of pHis and other phosphorylated amino acids in a cell lysate using MS is problematic. Depending on the precise conditions used in the MS analysis, there is a tendency for the phosphoryl group either to be lost before detection of the phosphorylated peptide or for it to transfer to another amino acid residue giving rise to false positives.^17–20^ In addition, as with other amino acid analysis methods such as Edman degradation, hydrolysis, and MS, samples are subjected to many potentially destructive and prolonged downstream steps which use acidic conditions either in sample preparation or in analysis. For example, in MS analysis of peptides, samples typically undergo overnight trypsinization (37 °C, pH 8), desalting (acidic pH) and HPLC (acidic pH) before analysis.^21^ Such conditions decompose acid labile phosphorylated amino acids such as pHis.^2, 22^

In this study, ^31^P NMR spectroscopy is introduced as a simple and direct detection method in the absolute quantification of pHis (and other phosphorylated amino acids) in a mammalian whole cell lysate, using the human bronchial epithelial cell line (16HBE14o- or HBE). The method is simple and involves minimal sample handling, and on average pHis was found to be much more abundant than previously thought: one third as abundant as pSer and pThr combined, strongly implying an important functional role for pHis in eukaryotic cells. The procedure is likely applicable for the absolute quantification of phosphorylated amino acids in other cells.

## Results

### Preliminary experiments

To address the question of how abundant pHis is in a mammalian cell using ^31^P NMR spectroscopy, it was first important to determine the regions in which pHis signals typically appear in a ^31^P NMR spectrum. τ-pHis and π-pHis were synthesized by the reaction of His with potassium phosphoramidate in water to allow determination of reference ^31^P chemical shifts of each isomer.^23^ Chemical shifts of - 4.99 ppm and - 5.76 ppm were observed for τ-pHis and π-pHis respectively (Supplementary Fig. S2a and b), entirely consistent with those previously reported.^23, 24^ To establish proof of concept and to identify ^31^P NMR chemical shifts characteristic of pHis in a protein environment under conditions that could be used to analyze proteins from a cell lysate, it was important to establish a protein standard. MS studies have shown that myoglobin can be phosphorylated selectively on His residues using potassium phosphoramidate in water to give myoglobin-pHis (Myo-pHis).^12, 25^ Potassium phosphoramidate was used to phosphorylate myoglobin in water, at 25 °C for 15 h.^25^ pHis is more stable under basic conditions (pH 10-12), and unstable under neutral and acidic conditions,^26^ and therefore Myo-pHis was buffer exchanged into 0.1 M Na_2_CO_3_/NaHCO_3_, at pH 10.8 (which also removed any unreacted potassium phosphoramidate), and subsequently concentrated and analyzed by ^31^P NMR spectroscopy. Multiple pHis residue signals were observed in the chemical shift range of - 4.40 to - 5.76 ppm (Fig. 1aii); additionally an inorganic phosphate (Pi) signal at 2.57 ppm and a signal at 8.27 ppm which was characteristic of pLys ^27^ (later confirmed by HMBC, see Fig. 1c) were also observed.

**Figure 1.**
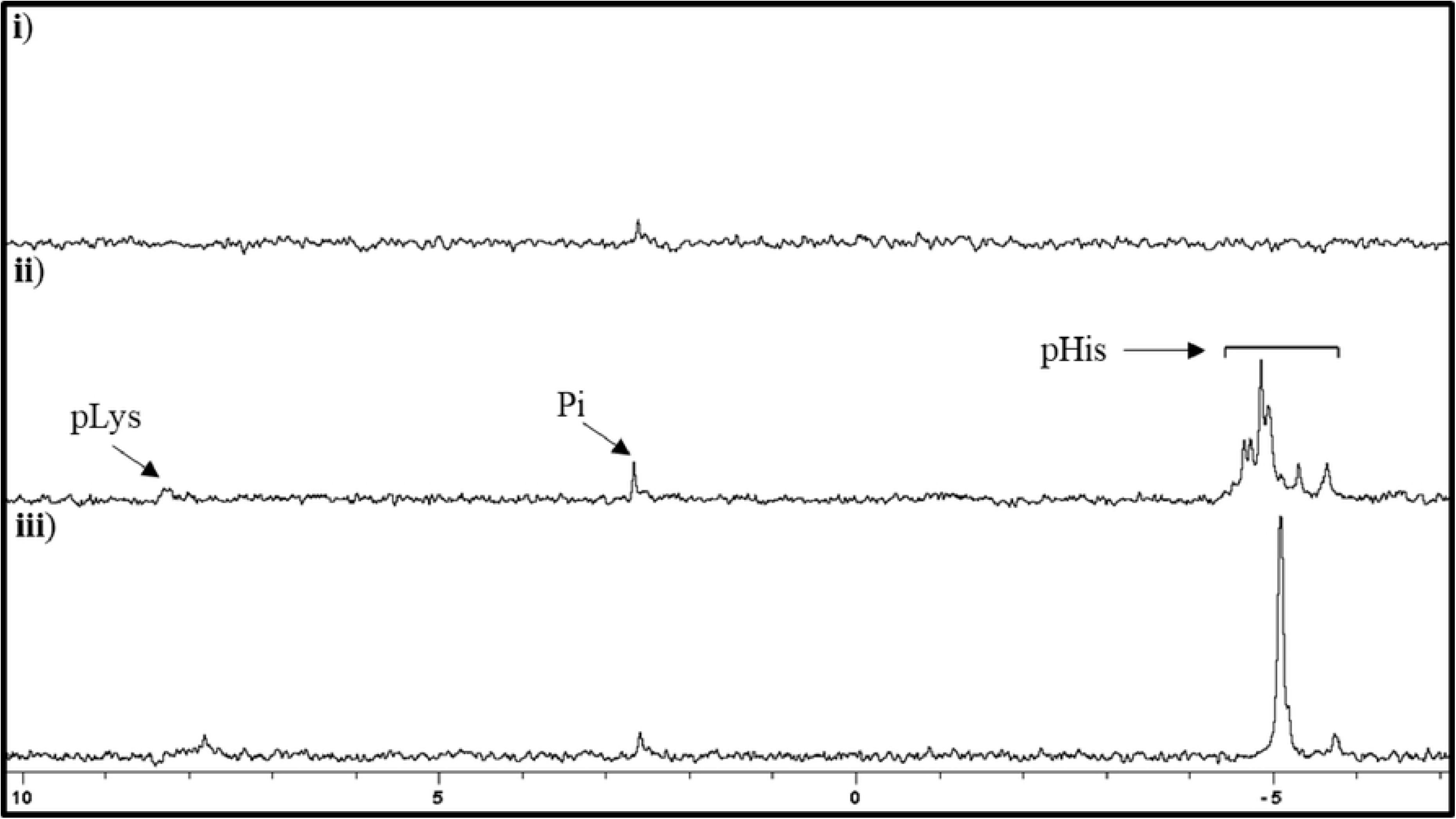
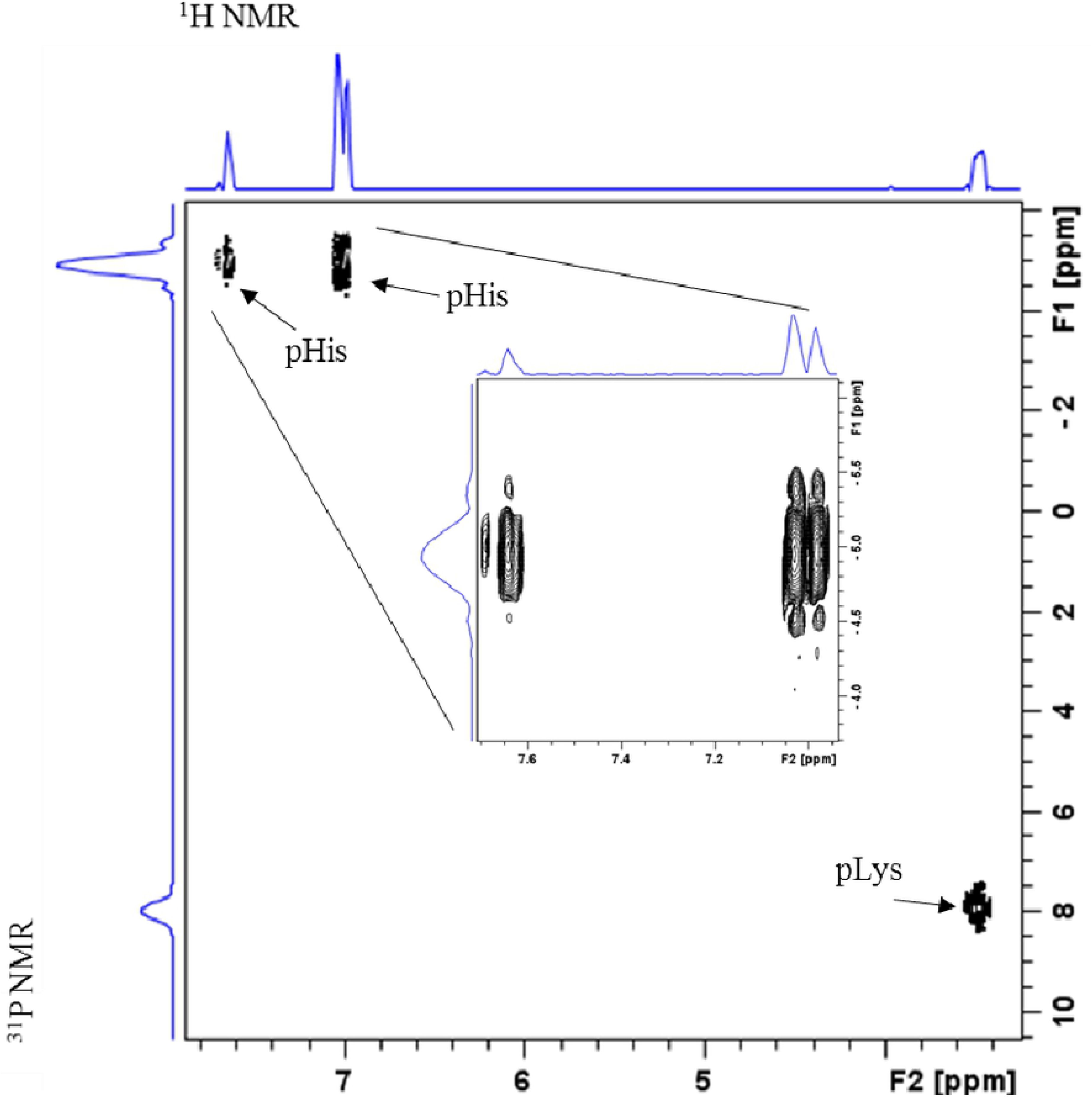
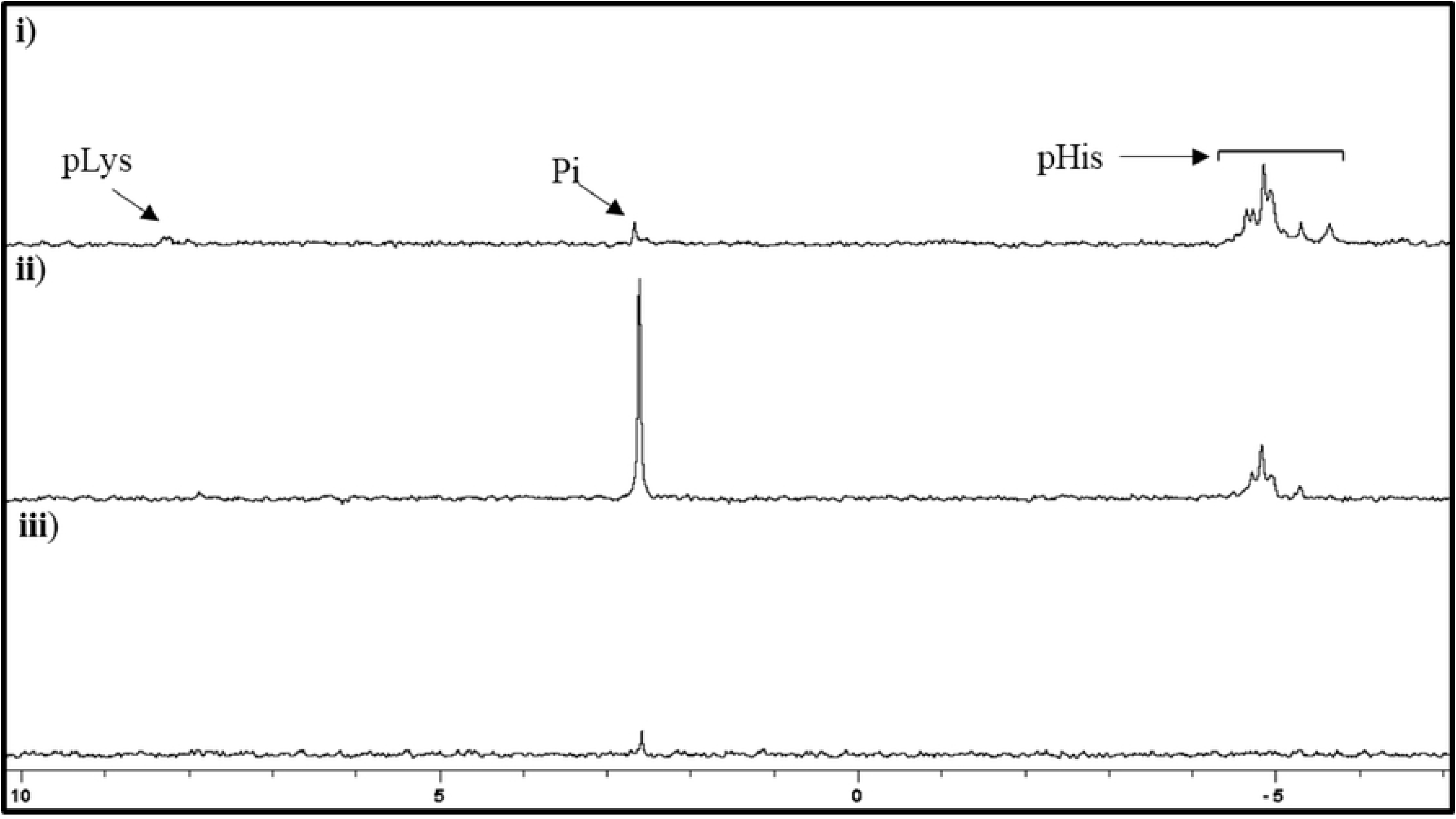
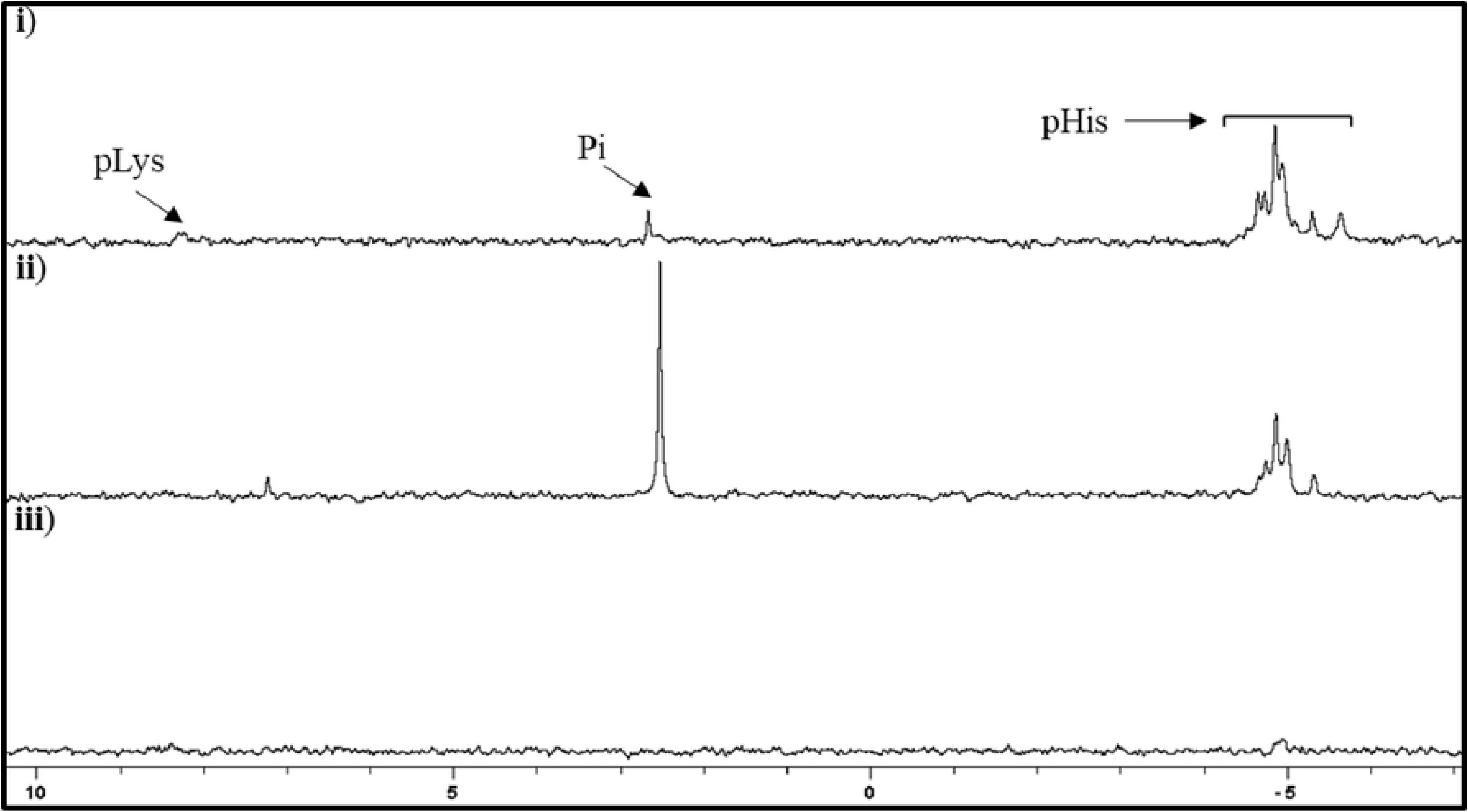
^31^P NMR spectra of Myo-pHis. **ai**) Myoglobin (17.7 mg/mL) in 91 mM Na_2_CO_3_/NaHCO_3_, 10 % (v/v) D_2_O, pH 10.8; **aii**) Myo-pHis (17.7 mg/mL) in 91 mM Na_2_CO_3_/NaHCO_3_, 10 % (v/v) D_2_O, pH 10.8. Myoglobin was phosphorylated with potassium phosphoramidate, was buffer exchanged into 0.1 M Na_2_CO_3_/NaHCO_3_, pH 10.8 and concentrated; **aiii**) Myo-pHis (13.8 mg/mL) in 71 mM Na_2_CO_3_/NaHCO_3_, 6.4 M urea, 10 % (v/v), D_2_O, pH 10.8. The Myo-pHis sample used was the same sample as Fig. 1aii. Note the Na_2_CO_3_/NaHCO_3_, and Myo-pHis concentration changed because of the addition of urea; **b**) HMBC NMR spectrum of Myo-pHis, sample Fig. 1aiii; **ci**) Myo-pHis (18.2 mg/mL) in 91 mM Na_2_CO_3_/NaHCO_3_, 10 % (v/v) D_2_O, pH 10.8, prepared as Fig. 1aii; **cii**) Myo-pHis after treatment with 1/50 (wt/wt) trypsin, at 37 °C for 16 h in 0.1 M NH_4_HCO_3_, pH 8.0. Na_2_CO_3_, NaHCO_3_ and D_2_O was subsequently added before ^31^P NMR spectroscopic analysis, **ciii**) Myo-pHis in 91 mM Na_2_CO_3_/NaHCO_3_, 10 % (v/v) D_2_O, pH 10.8 after trypsinisation and subsequent desalting; **di**) Myo-pHis (18.2 mg/mL) in 91 mM Na_2_ CO_3_ /NaHCO_3_, 10 % (v/v) D_2_O, pH 10.8, prepared as Fig. 1aii; **dii**) Myo-pHis after reduction with DTT (3 mmol), alkylation with IAA (14 mmol) and subsequent treatment with 2 % (wt/wt) trypsin, at 30 °C for 16 h in 0.1 M NH_4_HCO_3_, pH 8.0. Na_2_CO_3_, NaHCO_3_ and D_2_O was subsequently added before ^31^P NMR spectroscopic analysis; **diii**) Myo-pHis in 91 mM Na_2_CO_3_/NaHCO_3_, 10 % (v/v) D_2_O, pH 10.8, after trypsinization and subsequent desalting

**Figure 2.**
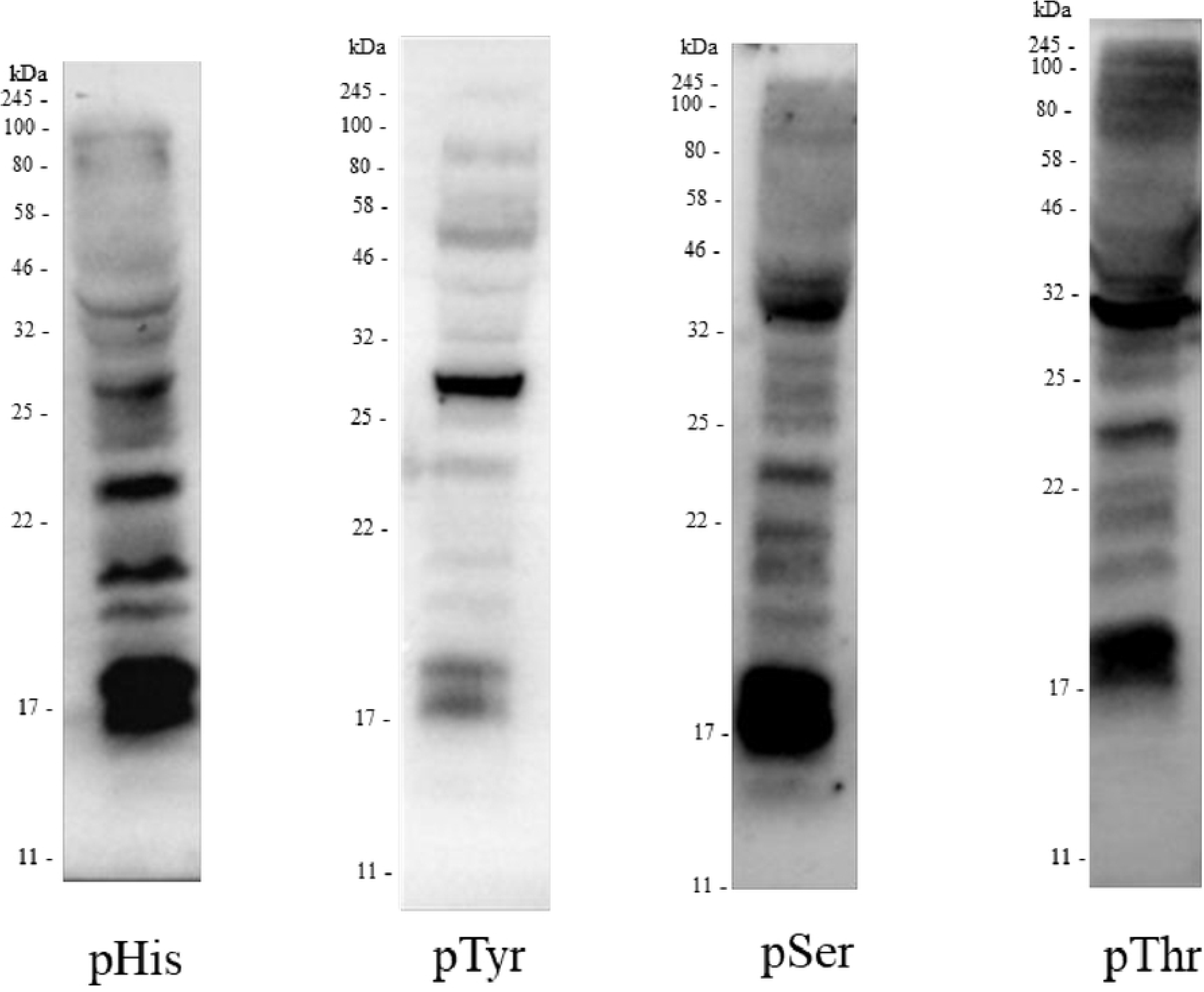
Western blot of 16HBE14o- cell lysates probed with phosphoantibodies. 16HBE14o- cell lysate (100 μg of protein) was probed with either pHis (ab2317090), pTyr (pY99), pThr, or pSer (Q5) antibody.

To reduce the tertiary structural effect of myoglobin on the pHis chemical shifts, Myo-pHis was treated with a denaturant, 7 M urea (Fig. 1aiii). Two distinct sets of signals in the ranges - 4.80 to - 5.38 ppm, and - 5.52 to - 5.98 ppm were observed in the ^31^P NMR spectrum of the denatured Myo-pHis. In heteronuclear multiple bond correlation (HMBC) NMR experiments of denatured Myo-pHis, cross peaks between ^1^H NMR aromatic protons and ^31^P NMR pHis residue signals are observed confirming phosphorylation had indeed occurred on His residues (Fig. 1b). On Western blots of Myo-pHis against commercially available pHis, pSer, pThr, and pTyr antibodies (Supplementary Fig. 3), no pSer, pThr, and pTyr residues were detected which is in agreement with the ^31^P NMR spectrum (Fig. 1) and reported MS data for Myo-pHis.^12, 25^ Furthermore, after acid treatment (~ pH 4) and heating (90 °C) of Myo-pHis, pHis and pLys residue signals were abolished, giving rise to an increased Pi signal suggesting the presence of acid labile residues (Supplementary Fig 4). pHis isomer distinction in a folded protein or peptide cannot always be made by comparing the pHis residue ^31^P NMR chemical shifts with pHis amino acid standards.^28^ However, comparison of the denatured Myo-pHis chemical shift regions of ~ - 4.94 ppm (major) and ~ - 5.61 ppm (minor) with the chemical shifts of τ-pHis and π-pHis amino acid standards (Supplementary Fig. S2a and b), suggests that the major pHis isomer in Myo-pHis was τ-pHis. τ-pHis is the more stable isomer of pHis^26^ and under conditions used to phosphorylate His, τ-pHis would be expected to be the major isomer present in Myo-pHis.

**Figure 3.**
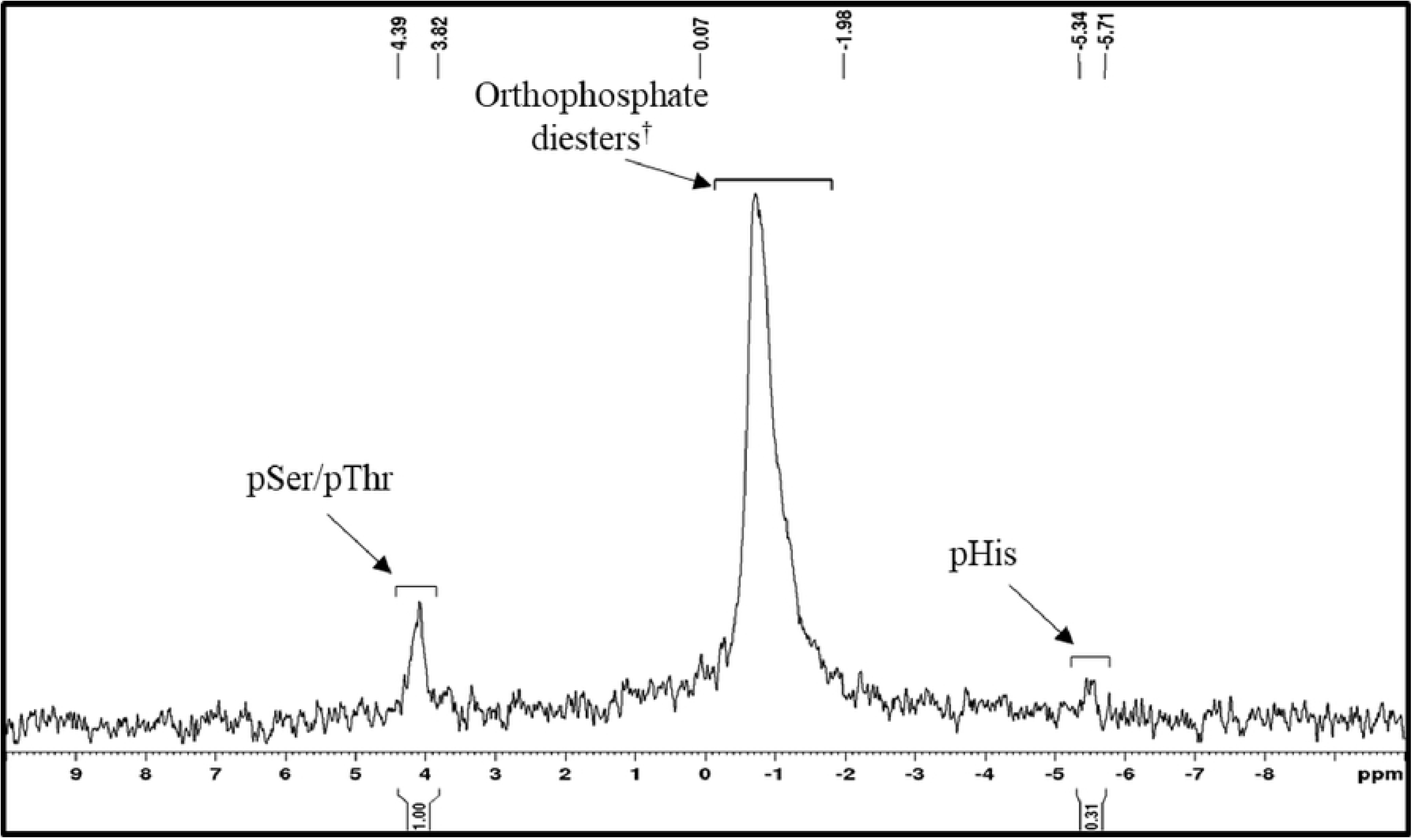
^31^P NMR spectrum of proteins from the 16HBE14o- cell lysate. Proteins from the 16HBE14o- cell lysate (15.6 mg/mL) in 91 mM Na_2_CO_3_/NaHCO_3_, 6.4 M urea, 10 % (v/v) D_2_O, pH 10.8. 16HBE14o- cells were lysed on ice in 0.1 M Na_2_CO_3_/NaHCO_3_, 7 M urea, pH 10.8. The sample was immediately sonicated, buffer exchanged into 0.1 M Na_2_CO_3_/NaHCO_3_, 7 M urea, pH 10.8, and concentrated. pSer and pThr signal assignments have been made using literature chemical shift values.^33, 34 †^DNA, RNA, phospholipids.^35^

**Figure 4.**
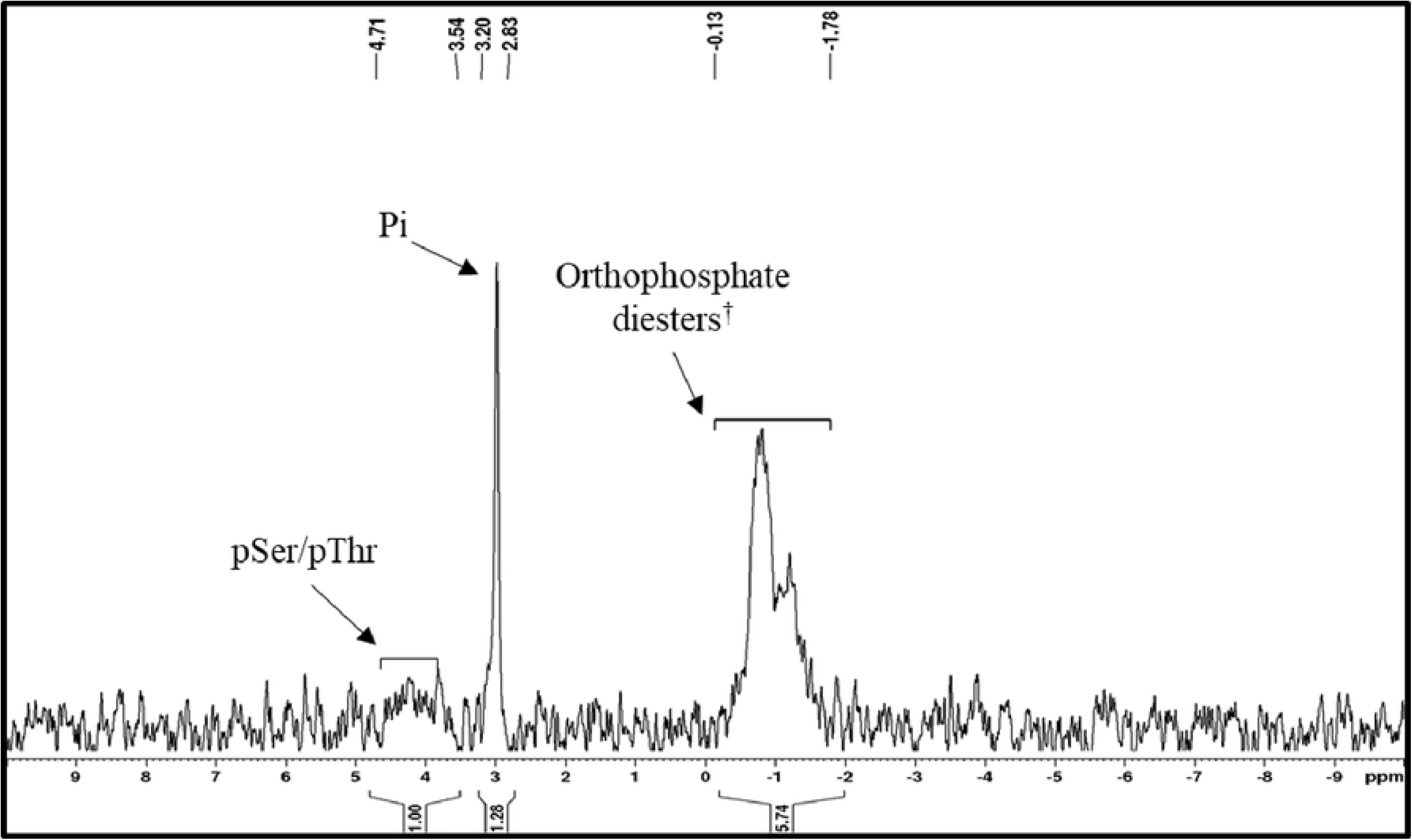
^31^P NMR spectra of proteins from 16HBE14o- cell lysate prepared by protein precipitation. **a**) Proteins from 16HBE14o- cell lysate in 91 mM Na_2_CO_3_/NaHCO_3_, 7.3 M urea, 10 % (v/v) D_2_O, pH 10.8. 16HBE14o- cells were lysed on ice in 30 mM Tris-HCl, 50 mM NaCl, complete protease inhibitor, pH 9.0.and the DNA was extracted following Antonioli *et al*.38 The precipitated protein was dissolved and buffer exchanged into 0.1 M Na_2_CO_3_/NaHCO_3_ 8 M urea, pH 10.8; **b**) Proteins from 16HBE14o- cell lysate in 91 mM Na_2_CO_3_/NaHCO_3_, 6.4 M urea, 10 % (v/v) D_2_O, pH 10.8. 16HBE14o- cells were lysed on ice in 0.1 M Na_2_CO_3_/NaHCO_3_, 7 M urea, pH 10.8, sonicated and the proteins were precipitated following Potel *et al*.^13^ pSer and pThr signal assignments have been made using literature chemical shift values.^33, 34 †^DNA, RNA, phospholipids.^35^

**Figure 5.**
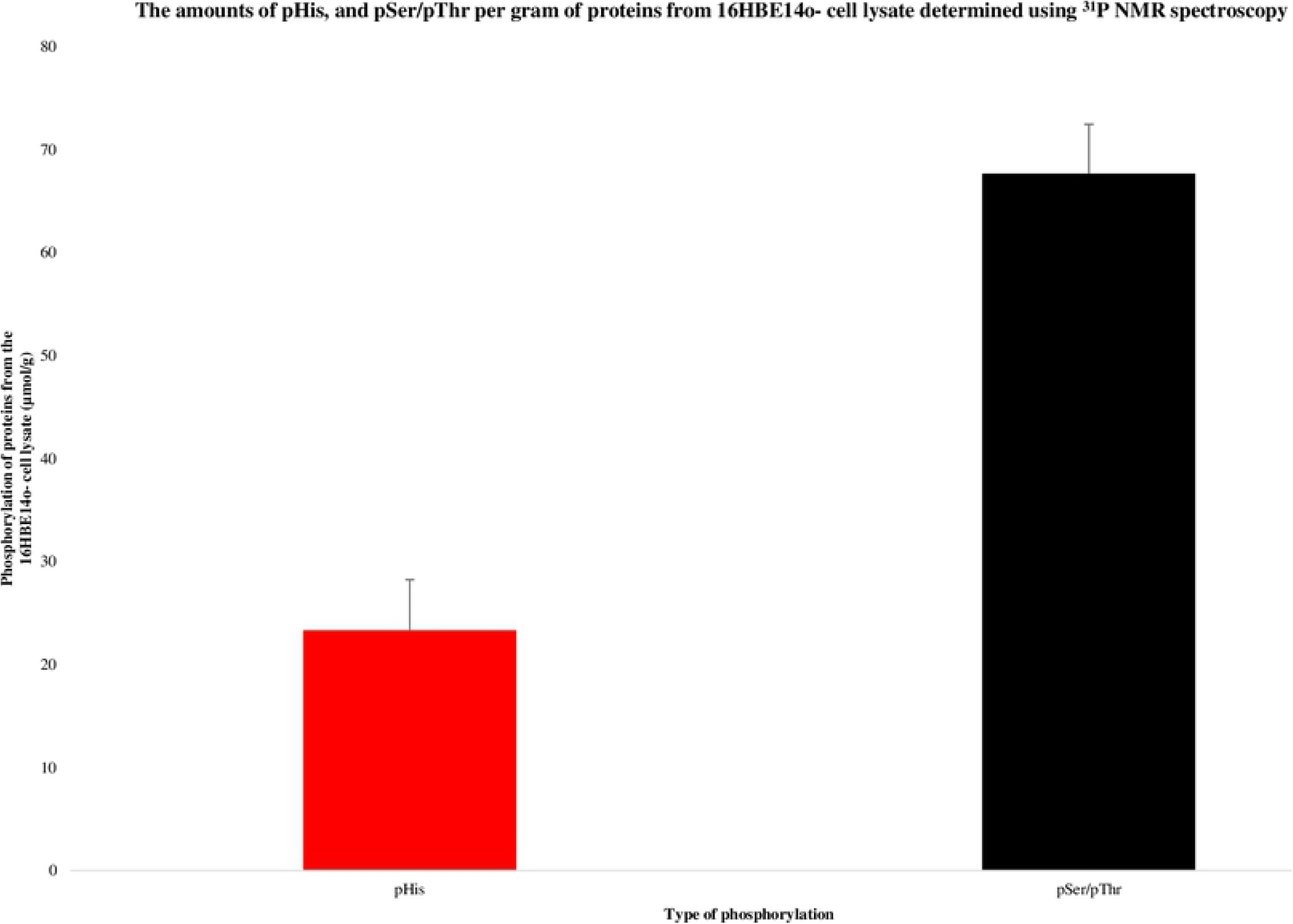
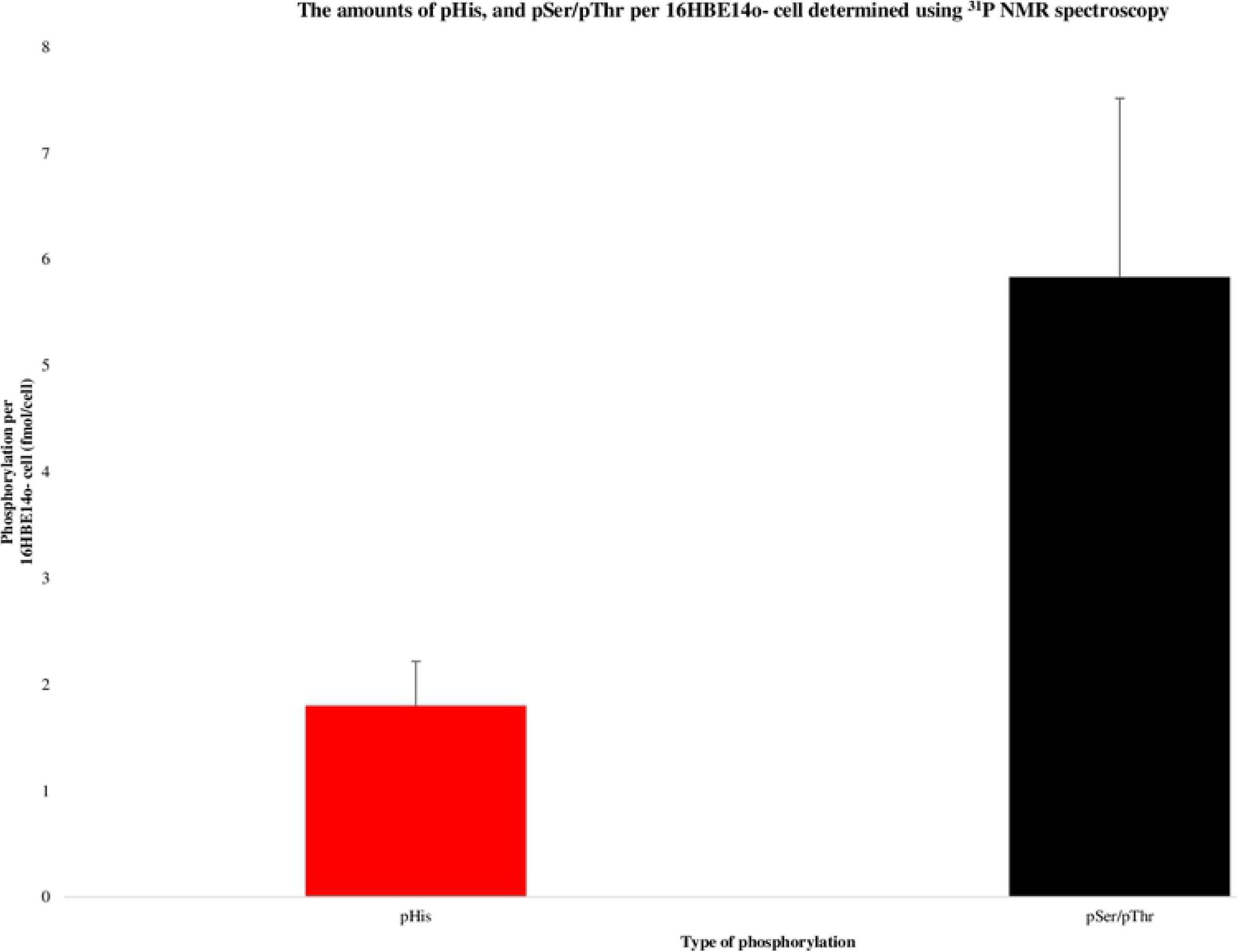
Quantification of pHis, pSer/pThr and polyphosphate using ^31^P NMR spectroscopy. **a)** The amounts of pHis, and pSer/pThr per gram of protein from 16HBE14o- cell lysate determined using ^31^P NMR spectroscopy; error bars show the standard error of the mean (*n* = 3); **b**) The amounts of pHis, and pSer/pThr per 16HBE14o- cell determined using ^31^P NMR spectroscopy; error bars show the standard error of the mean (*n* = 3).

To determine the number of pHis residues in the sample, a capillary tube containing an external standard (triphenylphosphine oxide, 10 mol % Cr(acac)_3_ in CDCl_3_) was used as an insert into the NMR tube containing Myo-pHis. Triphenylphosphine oxide was chosen as the external standard because it is a stable compound with a chemical shift ~ 30 ppm away from the pHis chemical shift region (Supplementary Fig. S5). The average number of pHis residues per myoglobin protein was determined to be 5.6 and the amount of pHis residues per gram of myoglobin was determined to be 0.32 mmol/g (Supplementary Fig. S5 for calculations). There are eleven His residues in myoglobin, all of which have been observed by MS to be phosphorylated by potassium phosphoramidate,^25^ and the quantitation here would suggest on average approximately half of His residues are phosphorylated. However, potassium phosphoramidate can phosphorylate both His imidazole nitrogens, and this possibility cannot be excluded for Myo-pHis.^23^

Trypsinization and desalting are typical steps used to process proteins from a cell lysate before MS analysis, and the conditions used are likely to result in the destruction of some pHis. To assess the impact of trypsinization conditions, Myo-pHis was trypsinized using typical conditions (37 °C, pH 8.0, 16 h)^21^ and then analyzed by ^31^P NMR spectroscopy (Fig. 1c). There was a significant change in the integration ratio of Pi and pHis residue signals from 1:15.8 for Myo-pHis (Fig. 1ci), to 1:0.97 for trypsinised Myo-pHis (Fig. 1ci). In addition, pLys residue signals also decreased. These results show extensive hydrolysis of both pHis and pLys residues under typical trypsinization conditions. The trypsin treated Myo-pHis (Fig. 1cii) was subsequently desalted using C-18 resin following typical acidic conditions. In the ^31^P NMR spectrum of the desalted sample no pHis residue signals were observed suggesting desalting also has a destructive effect on pHis residues (Figure 1ciii).^22^ Hardman *et al*. have detected Myo-pHis peptides by MS using milder trypsinization (Myo-pHis was first treated with dithiothreitol, and then iodoacetamide before the addition of trypsin at 30 °C, pH 8.0, 16 h) and desalting conditions (neutral solutions).^12^ Under these trypsinization conditions, the ^31^P NMR spectra (Fig. 1di and dii) integration ratios of Pi and pHis residue signals changed from 1:15.8 for Myo-pHis to 1:1.35 for trypsinized Myo-pHis showing a modest reduction in the amount of pHis residues that were hydrolyzed compared to typical trypsinization condition (*vide supra*). Following Hardman *et al*., Myo-pHis treated with trypsin was desalted using a C-18 resin and neutral solutions. The sample was subsequently analyzed by ^31^P NMR spectroscopy (Fig. 1diii). Relatively small pHis residue signals were observed when compared to the spectrum of Myo-pHis treated with trypsin (Fig. 1dii). The ^31^P NMR data shows that even using milder trypsinization and desalting conditions there was a significant loss of pHis.

### Quantitation of pHis in mammalian cells

The quantitative ^31^P NMR spectroscopic analysis of pHis residues developed on Myo-pHis was then applied to a more complex sample, namely the proteins from a cell lysate. Airway epithelia express His phosphorylated proteins and the 16HBE14o- cell line is a well-known airway epithelial cell line.^29, 30^ A Western blot of 16HBE14o- cell lysate using a pHis, antibody (ab2317090) detected several pHis bands confirming the presence of pHis residues (Fig. 2). Western blot and cell lysis procedures used pHis stabilizing conditions (see Methods). The pHis antibody selectivity was validated by acid treatment of the 16HBE14o- cell lysate which reduced the pHis signal, while base treatment retained the pHis signal (Supplementary Fig. S6). Similarly, Western blots of 16HBE14o- using pTyr, pThr, and pSer antibodies confirmed the presence of pTyr, pThr and pSer residues (Fig. 2).

To analyze the ^31^P NMR signals (specifically arising from pHis residues) of phosphoproteins from 16HBE14o- cell lysates, it was important to use a procedure that removed other phosphorus containing molecules (e.g. nucleoside triphosphates, oligonucleotides etc.) and as much DNA/RNA as possible, which could otherwise potentially dominate the ^31^P NMR spectrum. It was also important to keep the solution alkaline at all times, because pHis is degraded rapidly in acidic or neutral conditions. The preferred buffer was a weak base (such as Na_2_CO_3_/NaHCO_3_, pH 10.8, *vide supra*) because pSer and pThr are known to decompose by β-elimination on overnight treatment with a strong base (1 M NaOH, 37 °C).^2, 31^ Under these conditions, both pSer and pThr were found to be stable for more than two days (Supplementary Fig. S7a and b). Therefore, a robust, rapid and simple procedure was developed for preserving the native phosphorylation states.

Lysis buffers containing urea as the denaturant have been used to prepare cell lysates for MS analysis of pHis.^12, 13, 32^ Thus, 16HBE14o- cells were lysed on culture plates using 0.1 M Na_2_CO_3_/NaHCO_3_, 7 M urea, pH 10.8 and were immediately subjected to sonication. Sonication breaks DNA into small pieces which are subsequently removed along with other phosphorus containing small molecules during buffer exchange (Supplementary Fig. S8), while the denaturant ensures that there is reduced or no enzymatic activity. Using this procedure, pHis and pSer/pThr^33, 34^ residue signals in the ^31^P NMR spectrum were reproducibly observed (Fig. 3) Additional chemical shift signals from −0.05 to −1.70 ppm were also observed in Fig 3. The chemical shift regions in which these signals are present are characteristic of orthophosphate diesters,^35^ and thus have been assigned accordingly. In the basal state, the expected abundance of pTyr residues in 16HBE14o- cells is low^36^ and observable ^31^P NMR pTyr residue signals were not expected. Under the conditions used, pTyr is stable^37^ and is expected to have a chemical shift around 0.29 ppm (Supplementary Fig. S2e), but could not be assigned because any pTyr residue signals would overlap with signals due to orthophosphates.

An alternative DNA removal strategy described by Antonioli *et al*. that uses basic conditions to extract DNA, but precipitates the cell lysate proteins rather than keeping them in solution, was explored.^38^ Application of this method to the 16HBE14o- cell lysate (lysed in 30 mM Tris-HCl, 50 mM NaCl, complete protease inhibitor, pH 9.0) and subsequent ^31^P NMR spectroscopic analysis showed signals characteristic of orthophosphate diesters and pSer/pThr residues but no signals characteristic of pHis residues (Fig. 4a). Using protein precipitation conditions which have been previously used in the analysis of pHis by MS gave similar results (Fig 4b).^13^ Thus, it appears that protein precipitation is not beneficial in the analysis of pHis proteins from a cell lysate by ^31^P NMR spectroscopy.

In order to explore the wider applicability of this method, Jurkat cells were analyzed by Western blots and ^31^P NMR spectroscopy. Western blots of Jurkat cell lysate confirmed the presence of pHis, pTyr, pSer and pThr (Supplementary Fig. S9a). However, unlike the ^31^P NMR spectrum of the proteins from the 16HBE14o- cell lysate, there were no observable pHis residue signals in the ^31^P NMR spectrum of the proteins from the Jurkat cell lysate (Supplementary Fig. S9b) but there were signals for pSer/pThr residues. Proteins from both the Jurkat and 16HBE14o- cell lysate were lysed, and buffer exchanged identically. Since the pHis residue signals are only observed in the ^31^P NMR spectrum of the proteins from the 16HBE14o- cell lysate, this suggests that the abundance of pHis at the basal state can vary between different cell types, and that there may be different functional roles of pHis in different cell types.

For additional validation, the proteins from the 16HBE14o- cell lysate analyzed in Fig. 4a were chemically phosphorylated with potassium phosphoramidate which gave rise to ^31^P NMR signals from - 4.68 to - 6.13 ppm (Supplementary Fig. S10), confirming the signals observed in Fig. 3 are indeed pHis residues. Furthermore, as observed with Myo-pHis, acid treatment (~ pH 4) and heating (90 °C) of the proteins from the 16HBE14o- cell lysate resulted in loss of the pHis residue signals whilst giving rise to a Pi signal (Supplementary Fig. S11). Previously, relative phosphorylated amino acid abundances have been reported for proteins from a cell sample either as a percentage or ratio.^39^ However, since absolute quantitation is possible with ^31^P NMR spectroscopy (see *Preliminary experiments*), it is possible to determine the absolute amounts of different phosphorylated amino acid residues, as well as the relative amounts. In a triplicate ^31^P NMR spectroscopy analysis, pHis:pSer/pThr residue signal integration ratio was found to be on average 0.34:1 (Supplementary Fig. S12). Using an external standard, the average amounts of pHis, and pSer/pThr residues per mass of protein cell lysate were determined to be 23 μmol/g, and 68 μmol/g respectively (Supplementary Table S1). The amounts of pHis, and pSer/Thr residues per cell were determined to be 1.8 fmol/cell, and 5.8 fmol/cell respectively (Supplementary Table S2).

As observed with the Myo-pHis standard, the intensities of the ^31^P NMR signals assigned to pHis residues were found to reduce upon trypsinization (Fig. 6a). However, unlike Myo-pHis, new signals from 19.06 to 20.38 ppm (phosphonate region of the spectrum) in the ^31^P NMR spectrum of trypsinised proteins from the 16HBE14o- cell lysate were observed but no Pi signal. Phosphonates contain a phosphorus-carbon bond and such bonds are likely to arise from the reaction of a nucleophilic carbon species and phosphate electrophile. However, the mechanism by which these phosphonates arise and their identity is unclear but previous studies have observed phosphonate lipids giving similar chemical shift values.^40^ Since high urea concentration reduces trypsin activity,^21^ the 16HBE14o- cell lysates were buffer exchanged in preparation for trypsinization. It is likely during this step that enzymatic activity in the 16HBE14o- cell lysates was restored to some extent. Whether the emergence of the phosphonate signals and absence of Pi signal was due to a chemical or enzymatic transfer of the phosphoryl group is again unclear. In-solution-trypsinization for MS sample preparations generally use cell lysates that have not been dialyzed but have undergone reduction and alkylation under denaturing conditions, after which the denaturant is diluted, and the sample is treated with trypsin.^12, 41^ For simplicity, and to minimize sample handling, reduction and alkylation was not carried out. After desalting, the phosphonate signals were absent, which suggests that they are either labile under the desalting conditions used or have been removed during desalting. Noticeably, pSer/pThr residue signals appeared to have gradually increased after trypsinization and desalting, and the orthophosphate diester signals have decreased. The presence of the orthophosphate diester signals after the desalt suggests these orthophosphate diesters are covalently linked to proteins, which is not uncommon.^42^ For comparison purposes (see *Preliminary results*), proteins from the 16HBE14o- cell lysate were trypsinized and desalted following conditions by Hardman *et al.,* which included reduction and alkylation steps. A ^31^P NMR spectrum similar to Fig. 6iii was observed (Supplementary Fig. S14).^12^

**Figure 6.**
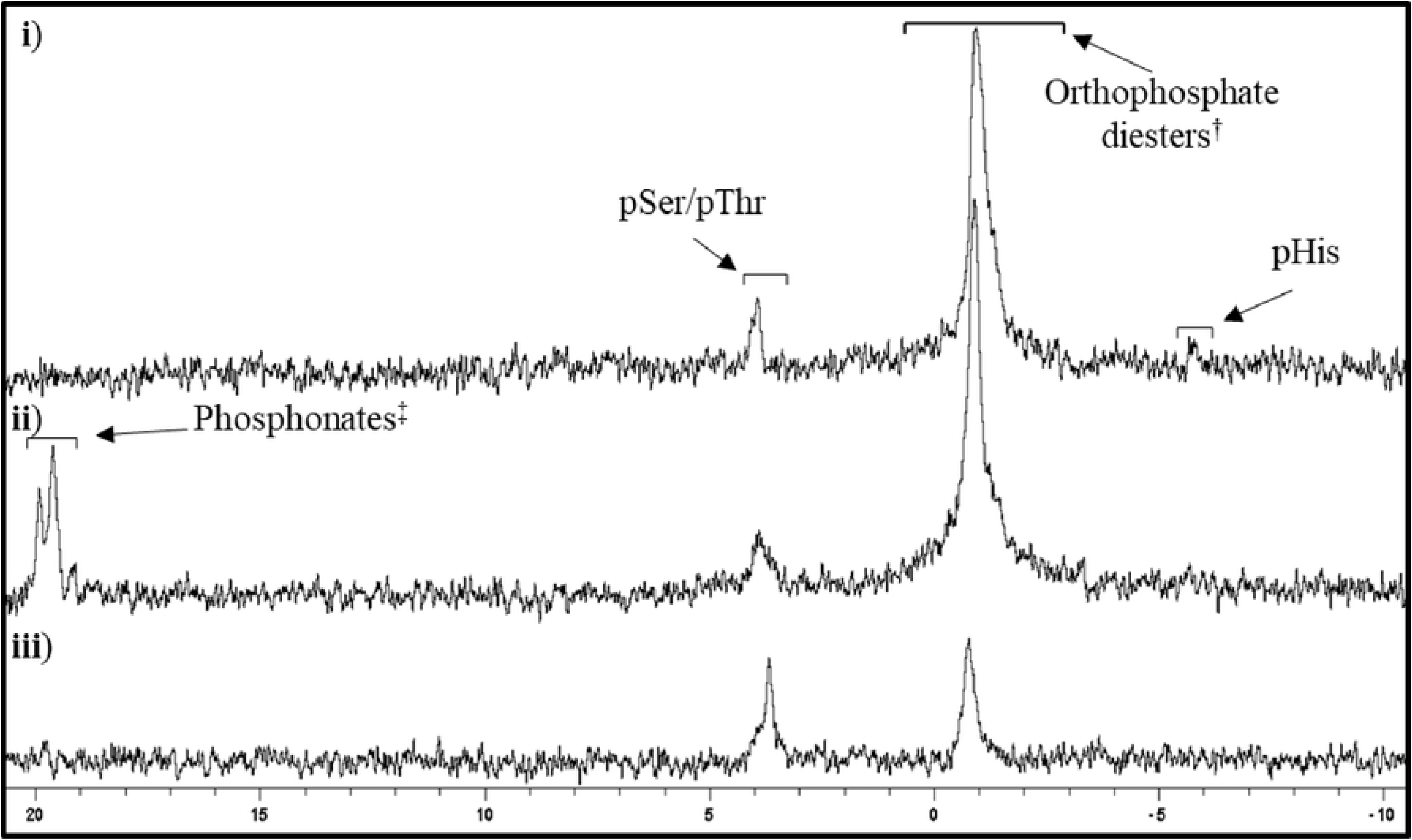
^31^P NMR spectra of proteins from 16HBE14o- cell lysate after trypsinization and desalting. **i**) Proteins from the 16HBE14o- cell lysate in 91 mM Na_2_CO_3_/NaHCO_3_, 6.4 M urea, 10 % (v/v) D_2_O, pH 10.8 prepared as Fig. 3; **ii**) sample Fig. 6i after treatment with 1/50 (wt/wt) trypsin, at 37 °C for 16 h in 0.1 M NH_4_HCO_3_, pH 8.0. Na_2_CO_3_, NaHCO_3_ and D_2_O was subsequently added before ^31^P NMR spectroscopic analysis; **iii**) sample Fig. 6ii in 91 mM Na_2_CO_3_/NaHCO_3_, 10 % (v/v) D_2_O, pH 10.8, after desalting.; Signal assignments have been made using literature chemical shift values.^33–35, 43 †^DNA, RNA, phospholipids.^35 ‡^New signals not present in Fig. 6i.

## Discussion

The study demonstrates unexpectedly high amounts of pHis in mammalian16HBE14o- cells. The study also suggests that pHis in cells may have previously gone undetected due to its labile nature under acidic and near neutral pH experimental conditions. By using a pHis stabilizing denaturing lysis buffer, and minimal sample processing steps followed by an upstream nondestructive analysis technique (i.e. ^31^P NMR spectroscopy), the amount of pHis residues was determined to be on average approximately one third the amount of pSer/pThr residues combined in proteins from the 16HBE14o- cell lysate. It is generally considered that the relative abundance of pSer, pThr and pTyr is approximately 83:15:1.5 in human cells,^44^ implying that pHis is likely to be about 20 times more abundant in human cells than pTyr (consistent with the results shown here), and that it is therefore likely to have hitherto unexplored functions. Finally, on subjecting proteins from the 16HBE14o- cell lysate to trypsinization, the ^31^P NMR pHis residue signals were abolished (Fig. 6) along with increased pSer/pThr residue signals, suggesting that some possible enzymatic or chemical transfer of phosphate from pHis or orthophosphate diesters to pSer/pThr may occur after cell lysis, providing a mechanism for potential false positives in phosphoproteomic experiments, especially under non-denaturing conditions.

The study demonstrates that ^31^P NMR spectroscopy can be applied to the absolute quantitation of pHis (and other phosphoamino acids) in a mammalian cell. The study also shows that ^31^P NMR spectroscopy can be used as a tool to monitor changes in phosphorylation resulting from subjecting a sample to different conditions. The simplicity of the procedure (only three sample processing steps before spectroscopic analysis: lysis, sonication, and simultaneous buffer exchange/concentration) highlights the potential broad applicability of the method in phosphoamino acid residue analyses of various cells/tissues or even small multicellular organisms. Since absolute quantitation is possible with ^31^P NMR spectroscopy, phosphorylation can be expressed in moles per gram of protein cell lysate or per cell, thereby allowing direct comparison between different cells or cell types.

Importantly, subjecting pHis-containing samples to trypsinization and desalting results in loss of pHis which has potential implication for MS sample preparations (Fig. 1 and 6). The artifactual increase of pSer/pThr following sample manipulation in cell lysates, shown in Fig. 6, is an interesting and surprising finding and suggests chemical or enzymatic transfer of phosphorus species. As such, it may be possible that the basal state abundances of pSer and pThr in cells may be significantly lower than currently recorded. Though the precise mechanism and importance of pHis phosphoryl transfer to serine (Ser) or threonine (Thr) residues are presently unclear, observations exist in the literature of *in vitro* Ser phosphorylation of kinase suppressor of Ras (KSR) in the presence of His kinase via nm23/NDPK, suggesting a potential His-to-Ser protein kinase pathway.^45^

Nevertheless, using a pHis stabilizing denaturing lysis buffer, the ^31^P NMR spectrum in Fig. 3 shows that, at the basal state, pHis residues are more abundant in mammalian 16HBE14o- cells than previously thought. Until now Ser, Thr and Tyr phosphorylated amino acid residues in mammalian cells have been thought to be the major players in cell signaling.

However, pHis may also be a major player in cell signaling. And though pHis is known to be involved in bacterial cell signaling,^2^ the roles of pHis in mammalian cells, and pHis proteins, their function, and associated (potential) kinases and phosphatases remain largely unknown.^14^ Genetic approaches in prokaryotes and lower eukaryotes revealed pHis residues are important in two-component signaling systems.^9^ The absence of corresponding vertebrate homologs of the two-component signaling systems led to the notion that perhaps pHis is predominantly a non-vertebrate cellular and signaling intermediate. This notwithstanding, a full characterization of pHis function in prokaryotes and lower eukaryotes also remains to be done; for example, an accurate abundance of pHis in non-vertebrate cells is still unclear. Hitherto, there was no procedure or method to provide practical insight to such key questions. One can speculate that pHis abundance relates to important mammalian cell requirements and cellular processes including cell signaling, immune system, energy stores/metabolism/homeostasis, and gene transcription/ regulation. One example of the importance of pHis is illustrated by the need to relay nascent ATP away from the outer rim of the mitochondrion to avoid negative charge build up such that energy production is compromised if phosphorylated histidine is unavailable.^46^ For example, mutation of NME3, a member of the NDPK family, underlies a fatal mitochondrial disorder,^47^ and double deletion of murine NME1 and NME2 is incompatible with postnatal life.^48^ It is possible that the high levels of pHis seen here are merely a consequence of the high levels of histidine and phosphate carriers in the cell, and have no biological implications. However, Nature tends to make use of such chemical opportunities, as witnessed in bacterial two-component signaling systems; and the observation of specific histidine-targeted phosphotransferases^15^ strongly implies a functional role.

In conclusion, this study introduces a ^31^P NMR spectroscopy method for absolute quantification of pHis and pSer/pThr in mammalian cells, thereby in principle allowing direct comparison between different cell types. This simple, direct method has allowed a longstanding question about the likely importance of pHis in mammalian cell biology to be addressed by demonstrating for the first time that pHis is a relatively abundant phosphorylation PTM in 16HBE14o- cells under basal conditions.

## Methods

### Quantitative ^31^P NMR spectroscopy of proteins

Norell® Standard Series™ 5 mm NMR tubes length 7 inches, inner diameter 4.2 mm were used. ^31^P spectra were acquired on a Bruker AVANCE III or Bruker AVANCE III HD spectrometer at 162.0MHz or 202.5MHz respectively. Data were acquired using an acquisition window of 9.69 kHz or 12.1 kHz respectively (59.9 ppm) with 32 k acquisition points (14-15 hours acquisition time), a 30° pulse and a relaxation delay of 8s. This pulse angle and recycle delay combination had previously been determined to be sufficiently long for complete relaxation, thus quantitative for all ^31^P containing species in the samples, including those with external standard which was doped with chromium(III) acetylacetonate to aid relaxation.

### External standard

Norell® high throughput 3mm NMR sampling tubes length 203 mm, inner diameter 2.41 mm were used as the external standard capillary. The capillary contained 2 or 4 mM triphenylphosphine oxide, 10 mol % chromium(III) acetylacetonate, dissolved in 150 μL of deuterated chloroform (CDCl_3_).

### ^1^H-^31^P HMBC spectroscopy

^1^H-^31^P HMBC spectra were acquired on a Bruker AVANCE III 400 spectrometer, using 1664 scans for each of 128 increments over an acquisition window of 9.7 kHz and 3.6 kHz (2 k points) in F1 and F2 respectively and optimized for a long-range coupling constant of 10 Hz.^24, 27^

### Materials and general experimental

All reagents used were purchased from Fluorochem, Millipore, Sigma, Alpha Aeser, VWR International, Thermo Fisher Scientific, Qiagen, and Bio-Rad Laboratories, Inc. Ultrapure water (18 MΩ) was used to make buffers and solutions unless stated otherwise. All solvents were of HPLC grade. ULTRA PURE AccuGel™ 29:1, 30 (w/v) 29:1 acrylamide: bis-acrylamide solution (gas stabilised) was used to make SDS-PAGE gels. PBS refers to phosphate buffered saline: 1.8 mM monopotassium phosphate, 10 mM disodium phosphate, 2.7 mM potassium chloride, 137 mM sodium chloride, pH 7.4. Pierce™ BCA protein assay kit, and Bio-Rad Bradford protein assay was used to estimate proteins. Thermo Scientific SuperSignal™ West Pico PLUS chemiluminescent substrate was used to develop blotted membranes. Sigma-Aldrich Medium 199 with Earle′s salts, sodium bicarbonate, phenol red, and without L-glutamine was used to culture 16HBE14o-. Gibco RPMI medium 1640 (1X), without L-glutamine was used to culture Jurkat cells. Gibco low endotoxin heat-inactivated foetal bovine serum was used to supplement media. Sigma-Aldrich Penicillin-Streptomycin; 10,000 units penicillin/mL and 10 mg streptomycin/mL was used to supplement media.

Sigma-Aldrich 200 mM L-Glutamine solution was used to supplement media. Sigma-Aldrich trypsin from bovine pancreas was used to trypsinize proteins. Sigma Trypsin-EDTA solution (1 ×, sterile; sterile-filtered, BioReagent, suitable for cell culture, 0.5 g porcine trypsin and 0.2 g EDTA • 4Na per liter of Hanks′ Balanced Salt Solution with phenol red) was used to trypsinize adhered 16HBE14o- cells. Roche complete protease inhibitor cocktail was used. Sigma-Aldrich myoglobin from equine heart was used. Sigma-Aldrich Gelatin from cold water fish skin was used. New England Biolabs pre-stained protein standard, broad Range (11-190 kDa) was used. Abcam pHis (ab2317090) at 1/1000 (v/v) dilution, Invitrogen pTyr (pY99) at 1/2000 (v/v) antibody dilution, Qiagen pThr (Q7) 1/200-1/2000 (v/v) antibody dilution, and Qiagen pSer (Q5) at 1/200-1/2000 antibody dilution, phospho antibodies were used. Dako rabbit anti-goat at 1/2000 (v/v) dilution, Dako mouse anti-rabbit at 1/2000 (v/v) dilution, and Abcam goat anti-Mouse IgG H&L (ab97023) at 1/10000 (v/v) dilution, immunoglobulins/ horseradish peroxidase were used as the secondary antibodies. Millipore® PVDF membranes (0.45 μm) were used to blot proteins. Silica gel 40-60 μm from VWR International was used. Merck TLC Silica gel 60 F254 TLC plates were used, and compounds were visualized by UV light (254 nm), 5 % (wt/v) ninhydrin in methanol. Sartorius Vivaspin 20 (3 and 10 kDa MWCO) were used for dialysis and concentration of protein solutions. Western blotted membranes were visualised using a Bio-Rad ChemiDoc™ XRS+ with image Lab™ software. Infrared spectra were recorded using a Perkin Elmer Paragon 100 FTIR Spectrophotometer by attenuated total reflectance (ATR). Only selected peaks are reported, and the absorption maxima are given to the nearest wavenumber (cm^−1^). pSer, pThr, pTyr, π-pHis, τ-pHis, and inorganic phosphate NMR spectra were recorded at room temperature. ^1^H NMR spectra were recorded using a Bruker AVANCE 400 spectrometer operating at 400.13 MHz, or Bruker AVANCE III HD 400 spectrometer operating at 400.23 MHz. ^13^C NMR spectra were recorded using a Bruker AVANCE 400 spectrometer operating at 100.61 MHz, or Bruker AVANCE III HD 400 spectrometer operating at 100.64 MHz. ^31^P NMR spectra were recorded using a Bruker AVANCE III HD 400 spectrometer operating at 162.02 MHz. Chemical shifts were measured relative to the residual solvent and expressed in parts per million (δ). The multiplicities are defined as s = singlet, d = doublet, t = triplet, q= quartet, quint. = quintet, sex. = sextet, m = multiplet, br = broad.. High-resolution mass spectra were measured using an Agilent Technologies 653 Accurate-Mass Q-TOF LC/MS operating in electrospray mode.

### Synthesis of amino(potassiooxy)phosphinic acid (Potassium phosphoramidate)

To a stirred mixture of water (100 mL), and 32 % (wt/wt) ammonia (50 mL), ice cold phosphorus(V) oxychloride (15.1 g, 9.2 mL, 98.5 mmol, 1.0 equiv.) was added dropwise at 4 °C, over 25 minutes. Effervescence was observed and the solution was stirred for a further 15 minutes. Acetone (500 mL) was added and the mixture was vigorously stirred for 5 minutes. The aqueous layer was partitioned and acidified with glacial acetic acid to pH 6. A precipitate was observed. After keeping the suspension at −20 °C for 60 minutes, the precipitate was filtered off and washed sequentially with ethanol (20 mL) and diethyl ether (20 mL). The air-dried crystals were added in small portions to 50 % (wt/v) potassium hydroxide (23 mL), and then heated to 60 °C for 20 minutes. The mixture was allowed to cool to room temperature and then acidified with glacial acetic acid to pH 6. The suspension was poured into ethanol (1150 mL) and left to stand at room temperature for 60 minutes. The precipitate was collected on a sintered funnel and washed sequentially with ethanol (2 × 30 mL) and diethyl ether (2 × 30 mL) and dried under reduced pressure. Potassium phosphoramidate (6.21 g, 47 %) white powder: **ν**_**max**_(thin film)/**cm**^**−1**^ 2849, 2473, 2175, 1614, 1462, 1413, 1150, 1073; ^**31**^**P NMR**(162 MHz, D_2_O) −3.33. The characterization data are comparable to the literature.^49^

### Synthesis of 2-amino-3-(3-phosphonoimidazol-4-yl)propanoic acid (π-pHis)

L-Histidine (0.251 g, 1.62 mmol, 1.0 equiv.), potassium phosphoramidate (0.57 g, 4.21 mmol, 2.6 equiv.) and water (4.5 mL) were stirred at 25 ° C for 40 minutes. π-pHis was chromatographically purified see (§) for detailed procedure. The combined product fractions were concentrated by rotary evaporation at 25 °C to approximately 4 mL whilst maintaining a pH between 10-12 using 2 M sodium hydroxide and 1 M hydrochloric acid. The remaining solution was aliquoted and snap frozen and stored at −80 °C. The concentration of π-pHis was determined to be 2.2 mg/mL by ^1^H NMR using 1,4-dioxane (0.001 g, 1 μL, 0.012 mmol) as the standard. π-pHis (0.0084 g, 2 % yield): *R*_F_ = 0.44 (65:8:22 EtOH/32 % (wt/wt) NH_3_/H_2_O); ^**1**^**H NMR**(400 MHz) 2.96 (dd, *J* = 8.0, 15.5 Hz, 1H), 3.19 (dd, *J* = 5.0, 15.5 Hz, 1H), 3.84-3.90 (m, 1H), 6.72 (s, 1H), 7.73 (s, 1H); ^**13**^**C NMR**(101 MHz) 30.6 (s), 55.3 (s), 126.2 (d, *J* = 8.0 Hz), 130.4 (d, *J* = 3.5 Hz), 140.4 (d, *J* = 5.0 Hz), 181.9 (s); ^**31**^**P NMR**(162 MHz) - 5.61; **m**/**z**(ESI+): 236.0431 (MH+, 100 % C_6_H_11_N_3_O_5_P requires 236.0400). The characterization data are comparable to the literature.^23^

### Synthesis of 2-amino-3-(1-phosphonoimidazol-4-yl)propanoic acid (τ-pHis)

L-Histidine (0.251 g, 1.62 mmol, 1.0 equiv.), potassium phosphoramidate (0.57 g, 4.21 mmol, 2.6 equiv.) and water (4.5 mL) were stirred at 25 ° C for 16 hours. τ-pHis was chromatographically purified, see (§) for detailed procedure. The combined product fractions were concentrated by rotary evaporation at 25 °C to approximately 8 ml whilst maintaining a pH between 10-12 using 2 M sodium hydroxide and 1 M hydrochloric acid. The remaining solution was aliquoted and snap frozen and stored at −80 °C. The concentration of τ-pHis was determined to be 4.9 mg/mL by ^1^H NMR against 1,4-dioxane as the standard. τ-pHis (0.103 g, 27 % yield): *R*_F_ = 0.40 (65:8:22 EtOH/32 % (wt/wt) NH_3_/H_2_O); ^**1**^**H NMR**(400 MHz) 2.53 (dd, *J* = 9.0, 14.5 Hz, 1H), 2.79 (dd, *J* = 4.5, 14.5 Hz, 1H), 3.32-3.38 (m, 1H), 6.86 (s, 1H), 7.56 (s, 1H); ^**13**^**C NMR**(101 MHz) 32.6 (s), 56.0 (s), 117.8 (d, *J* = 5.5 Hz), 137.0 (d, *J* = 8.5 Hz), 139.0 (d, *J* = 5.0 Hz), 181.2 (s); ^**31**^**P NMR**(162 MHz) - 4.78; **m**/**z**(ESI+): 236.0431 (MH+, 100 % C_6_H_11_N_3_O_5_P requires 236.0400). The characterization data are similar to the literature.^23^

§ The following flash chromatography procedure was carried out at 4 °C and a positive pressure of 0.68 atmospheres. The sample was loaded on to a pre-equilibrated (85:4:1 ethanol/32 % (wt/wt) ammonia/water) silica gel column (4 cm diameter column with 27 cm of silica) and eluted with a mixture of ethanol, 32 % (wt/wt) ammonia and water gradient. (Initially 300 mL of an 85:4:1 mixture was added: once 150 mL was eluted, 150 mL of a 75:4:10 mixture was added. After a further 150 mL was eluted an additional 150 mL of this solvent was added. Subsequently the same cycle was repeated with solvent mixture ratio of 70:4:15 (300 mL), and 60:4:25, (300 mL) until the compound was eluted). The first 750 mL was run off and 15 mL fractions were collected. The product fractions were collected such that 5 fractions between the product and the first (for π-pHis) and last (π-pHis) co-eluted π- and τ-pHis mixture were discarded as analysed by TLC.

### Synthesis of phosphorylated myoglobin (Myo-pHis), and dialysis

Dialysis buffer: 0.1 M sodium carbonate/bicarbonate, pH 10.8.

Myoglobin from equine heart (50 mg, 2.8 μM, 1 equiv.) was dissolved in water (4.5 mL). Potassium phosphoramidate (0.61 g, 4.5 mmol, 1585 equiv.) was added and the mixture was stirred at 25 °C for 15 hours. The mixture was added to a Vivaspin 20 (10 kDa, MWCO) and concentrated down to ~1 mL by centrifugation (6 °C, 3220 RCF). The solution was made up to 5 mL with dialysis buffer and the solution was again concentrated down to ~1 mL as described above; this step was repeated 7 times. The sample was made up to 2 mL using dialysis buffer. The Myo-pHis protein concentration was determined by Bradford assay and myoglobin was used to generate the standard curve. 10 % (v/v) D_2_O was added to the Myo-pHis solution and analysed by ^31^P NMR spectroscopy. The sample was stable for weeks at 4 °C and could be snap frozen and stored at 80 °C (freeze thaw cycles were avoided). For ^31^P NMR spectroscopy of denatured Myo-pHis, urea (210 mg) was added to the Myo-pHis (500 μL) solution followed by D_2_O (50 μL) before analysis.

### Acid treatment of Myo-pHis

The pH of Myo-pHis was reduced to ~pH 4 using glacial acetic acid and it was then heated at 90 °C for 45 min. A precipitate was observed. Sodium carbonate (10 mg) and sodium hydrogen carbonate (1 mg) was added. To help dissolve the precipitate, urea (210 mg) and ethylenediaminetetraacetic acid disodium salt dihydrate (9 mg) were added and left overnight. 10 % (v/v) D_2_O was added to the sample before analysis by ^31^P NMR spectroscopy.

### Sodium Dodecyl Sulfate-Polyacrylamide Gel Electrophoresis (SDS-PAGE)

Resolving gel: 12 % (wt/v) acrylamide, 0.1 % (w/v) sodium dodecyl sulfate, 0.1 % (w/v) ammonium persulphate, 0.001 % (v/v) tetramethylenediamine, 0.38 M tris(hydroxymethyl)aminomethane-hydrochloride, pH 8.8.

Stacking gel: 4% (wt/v) acrylamide, 0.1 % (w/v) sodium dodecyl sulfate, 0.1 % (w/v) ammonium persulphate, 0.001 % (v/v) tetramethylenediamine, 0.125 Mtris(hydroxymethyl)aminomethane-hydrochloride, pH 8.8.

Freshly prepared resolving gel solution was immediately poured between 10 cm × 10 cm (1 mm spacers) and 10 cm × 10 cm (notched) glass plates. Isopropanol (~200 μL) was slowly added to the top. The gel was left to polymerise at room temperature for 30 minutes. The isopropanol was poured away, and the top of the gel was washed with water (× 3). Excess water was soaked up with a filter paper. Freshly prepared stacking gel (pH 8.8) solution was poured over the resolving gel to fill the cavity. A 10 well comb (~30 μL capacity per well) was inserted and the gel was allowed to form at room temperature for 2 hours. Gels were used immediately or stored in plastic with a damp (running buffer) tissue for up to 3 days.

### Culturing of human bronchial epithelial (16HBE14o- or HBE) cell line

Full serum medium: Medium 199, supplemented with 10 % (v/v) fetal bovine serum, 1.3 mM of L-glutamine, 80 μg/mL streptomycin, 80 units/mL penicillin.

16HBE14o- cell line was cultured in 57 cm^2^ dishes in full serum medium (10 mL) and incubated at 37 °C in a 5 % CO_2_ atmosphere. The medium was changed every 2-3 days by discarding the old medium, washing the cells with PBS (3 mL) and then replacing with fresh full serum medium (10 mL). Once the 16HBE14o**-**cells were between 70-90 % confluency, they were split or harvested.

### Culturing of Jurkat cell line

Full serum medium: RPMI medium 1640, supplemented with 10 % (v/v) fetal bovine serum, 1.3 mM of L-glutamine, 100 μg/mL streptomycin, 100 units/mL penicillin.

Jurkat cells were cultured in T75 flasks in full serum medium (30 mL) and incubated at 37 °C in a 5 % CO_2_ atmosphere. The cells were harvested or split 1:3 once cell densities of 1 × 10^6^ cell/mL were reached.

### Lysis of 16HBE14o- and Jurkat cells for Western blots

Western blot lysis buffer: 150 mM sodium chloride, 0.5 % (wt/v) sodium deoxycholate, 1 % (v/v) Triton X-100, 0.1 % (wt/v) sodium dodecyl sulfate, 10 mM sodium fluoride, 5 mM sodium orthovanadate, 10 mM sodium pyrophosphate, complete protease inhibitor, 50 mM tris(hydroxymethyl)aminomethane-hydrochloride, pH 8.8.

For preparation of 16HBE14o- cell lysate for Western blots, media from 57 cm^2^ cell culture dishes with adherent 16HBE14o- cells was discarded and the cells were washed with PBS (5 mL). The cell culture dish was cooled on ice and any residual PBS was discarded. Cells were scraped into ice cold lysis buffer (100 μL). After centrifugation (4 °C, 8950 RCF, 1 min), the supernatant was taken and used immediately or snap frozen and used within 3 days.

For preparation of Jurkat cells for Western blots, cell suspension (10-14 million cells total) were centrifuged (room temp., 453 RCF, 10 min). The supernatant was discarded, and the cells were suspended in PBS (5 mL). After centrifugation (room temp., 453 RCF, 10 min) the supernatant was discarded, and the cells were cooled on ice. Ice-cold lysis buffer (100 μL) was added and the mixture was agitated by vortexing before centrifugation (4 °C, 8950 RCF, 1 min). The supernatant was taken and used immediately or snap frozen and used within 3 days.

### Western blot

5× (concentrated) sample buffer: 10 % (wt/v) lithium dodecyl sulfate, 40 % (wt/v) glycerol, 0.02 % (wt/v) bromophenol blue, 50 mM ethylenediaminetetraacetic acid disodium salt dihydrate, 500 mM dithiothreitol, 300 mM tris(hydroxymethyl)aminomethane-hydrochloride, pH 9.0. 1× sample buffer was made by diluting 5× sample buffer in water.

Resolving buffer: 0.1 % (w/v) sodium dodecyl sulfate, 192 mM Glycine, 25 mM tris(hydroxymethyl)aminomethane, pH 8.3.

Transfer buffer: 192 mM glycine, 25 mM tris(hydroxymethyl)aminomethane, pH 8.3.

Wash buffer: 165 mM sodium chloride, 0.05 % (v/v) Tween 20, 10 mM tris(hydroxymethyl)aminomethane-hydrochloride, pH 8.0.

Blocking buffer: 0.2 % (v/v) gelatin from cold water fish skin, 165 mM sodium chloride, 0.05 % (v/v) Tween 20, 10 mM tris(hydroxymethyl)aminomethane-hydrochloride, pH 8.0.

Protein samples were treated with 5× sample buffer to give a 1× final concentration and then made up to 30 μL with 1× sample buffer. The samples were left at room temperature for 15 minutes and then loaded in to the SDS-PAGE gel wells. The samples were resolved using resolving buffer with the tank immersed in ice, at 120 V for the first 10 minutes and then 180 V for 50-60 minutes. The resolved proteins were immediately electro blotted onto a methanol activated PVDF membrane using transfer buffer with the tank immersed in ice at 100 V for 1 hour. The following steps were carried out at room temperature. Blocking buffer (75 mL) was added over the membrane (~8 × 8 cm) in a covered tray (10 cm × 14 cm) and left for 1 hour on an orbital shaker. The solution was discarded. The membrane and primary antibody diluted in blocking buffer (10 mL) was added to a centrifuge vial (50 mL) and left for 1 hour on a tube roller. The solution was discarded, and the membrane was put back in the tray on an orbital shaker and washed (6 × 5 min) with wash buffer (50 mL). The membrane and secondary antibody diluted in blocking buffer (10 mL) were added to a centrifuge vial (50 mL) and left for 1 hour on a tube roller. The solution was discarded, and the membrane was put back in the tray on an orbital shaker and washed (6 × 5 min) with wash buffer (50 mL). The membrane was then incubated in chemiluminescence solution for 1 minute. After draining off the excess chemiluminescence solution, the membrane was imaged.

### Trypsinization of Myo-pHis and desalting under typical conditions

The method followed Chen *et al*.^21^ with minor changes.

Dialysis buffer: 0.1 M ammonium bicarbonate, pH 8.0.

Myo-pHis (20 mg) was buffer exchanged (3 cycles of buffer exchange) into dialysis buffer using a Vivaspin 20 (3 kDa MWCO) and made up to 1 mL. 1/50 (wt/wt) trypsin was added and the mixture was shaken (600 rpm) at 37 ° C for 16 hours. To half of the mixture (~ 500 μL), sodium carbonate (10 mg), sodium hydrogen carbonate (1 mg), and D_2_O (50 μL) was added and analyzed by ^31^P NMR spectroscopy.

A C-18 resin Sep-Pak cartridge (130 mg) was conditioned sequentially with acetonitrile (2.6 mL), 50 % (v/v) acetonitrile, 0.1 % trifluoroacetic acid (2.6 mL) and 0.1 % trifluoroacetic acid (2.6 mL). The remaining sample (~500 μL) was made up to 2 mL using water. Half of the sample (1 mL) was acidified to pH 3 using aqueous 10 % (v/v) trifluoroacetic acid and passed through the cartridge. The flow through was collected and passed through the cartridge again; this step was repeated once more. The resin was washed with 0.1 % (v/v) trifluoroacetic acid (2.6 mL), and then extracted with 50 % (v/v) acetonitrile, 0.1 % (v/v) trifluoroacetic (0.6 mL). The other half of the sample (1 mL) was desalted in the same way using a fresh cartridge. The solvent from the combined fractions was removed under reduced pressure. The remaining residue was solubilized in 0.1 M sodium carbonate/hydrogen carbonate, pH 10.8 (500 μL), and analyzed by ^31^P NMR spectroscopy after the addition of D_2_O (50 μL).

### Trypsinization of Myo-pHis and desalting under milder conditions

The method followed Hardman *et al*.^12^ with minor changes.

Dialysis buffer: 0.1 M ammonium bicarbonate, pH 8.0.

Myo-pHis (20 mg) was buffer exchanged (3 cycles of buffer exchange) into dialysis buffer using a Vivaspin 20 (3 kDa MWCO) and made up to 1 mL. The sample was treated with dithiothreitol (7 mM) and shaken (600 rpm) at 30° C for 20 minutes. After allowing the mixture to cool to room temperature, iodoacetamide (14 mM) was added and the mixture was left in the dark at room temperature. Dithiothreitol (3 mM) was added followed by 2 % (wt/wt) trypsin and the mixture was shaken (600 rpm) at 30 ° C for 16 hours. To half of the mixture (~ 500 μL), sodium carbonate (10 mg), sodium hydrogen carbonate (1 mg), and D_2_O (50 μL) was added and analysed by ^31^P NMR spectroscopy.

A C-18 resin Sep-Pak cartridge (130 mg) was conditioned sequentially with methanol (2.6 mL), 50 % (v/v) acetonitrile (2.6 mL) and water (2.6 mL). The remaining sample (~500 μL) was made up to 2 mL using water and half of the sample (1 mL) was passed through the cartridge. The flow through was collected and passed through the cartridge again; this step was repeated once more. The resin was washed with water (2.6 mL), and then extracted with 50 % (v/v) acetonitrile (0.6 mL). The other half of the sample (1 mL) was desalted in the same way using a fresh cartridge. Both desalted samples were combined and solvent was removed under reduced pressure. The remaining residue was solubilized in 0.1 M sodium carbonate/hydrogen carbonate, pH 10.8 (500 μL), and analysed by ^31^P NMR spectroscopy after the addition of D_2_O (50 μL).

### Preparation of proteins from the 16HBE14o- and Jurkat cell lysate for ^31^P NMR spectroscopy

The lysis and dialysis procedure was done as quickly as possible.

Lysis buffers: 16HBE14o- cells were lysed in 0.1 M sodium carbonate/bicarbonate, 7 M urea, pH 10.8.

Full serum medium: Medium 199 (including phenol red), supplemented with 10 % (v/v) fetal bovine serum, 1.3 mM of L-glutamine, 80 μg/mL streptomycin, 80 units/mL penicillin.

Dialysis buffer: 7 M urea, 0.1 M sodium carbonate/bicarbonate, pH 10.8.

The medium was removed from five cell culture dishes (57 cm^2^) containing adhered 16HBE14o- cells between 70-90 % confluent. The medium was discarded, and the cells were washed with PBS (10 mL). The petri dishes were cooled on ice and care was taken to remove as much PBS as possible. Ice cold lysis buffer (150 μL) was added to each petri dish. Each petri dish was carefully shaken to evenly spread the lysis buffer. The cells were scraped into the lysis buffer and the lysate from each dish was transferred to another vial on ice. The lysis procedure was repeated three more times for a total of twenty cell culture dishes. For a cell count, five petri dishes were taken, and the cells were washed as described above. See (∥) below for cell count procedure. The combined cell lysate was sonicated on ice (45% amplitude 10 × 5 sec bursts, 59 sec rest between each burst). The lysate (~ 6 mL) was added to a Vivaspin 20 protein concentrator spin column (3 kDa MWCO) and concentrated down to ~1.5 mL by centrifugation (6 °C, 3220 RCF, ~2.5 hr). 1 mL dialysis buffer was added, and the mixture was concentrated down as above to ~ 1.5 mL. This step was repeated 14 times.

The sample finally was concentrated down to ~0.5 mL. 10 % (v/v) D_2_O was added to a portion of this sample which was analysed by ^31^P NMR spectroscopy (for quantitative analysis 300 μL was used to accommodate the capillary containing the external standard). The 16HBE14o- cell lysate protein concentration was quantified by BCA assay, using BSA to generate the standard curve. The sample was stable for weeks at 4 °C and could be snap frozen and stored at 80 °C (freeze thaw cycles were avoided).

(∥) Each dish was treated with 2 mL of trypsin, at 37 °C for 7 minutes. A cell scraper was used to help dislodge any weakly adhered cells. The cells in each petri dish were diluted with full serum medium to give a total volume of 10 mL and transferred to a separate vial. In a triplicate cell count, 10 μL was taken from each cell suspension to determine the average cell count using a hemocytometer. The average cell count of each cell suspension was then averaged between the five separate samples, to give an average cell count.

Jurkat cell (~240 × 10^6^ cells) were pelleted by centrifugation (room temp., 207 RCF, 10 min) and the supernatant was discarded. The cells were washed with PBS (10 mL) and the centrifugation cycle was repeated. The cells were lysed on ice using lysis buffer (6 mL) and prepared for ^31^P NMR analysis as described above.

### Trypsinization of proteins from 16HBE14o- cell lysate and desalting under typical conditions

The method followedChen *et al*.^21^ with minor changes.

Dialysis buffer: 0.1 M ammonium bicarbonate, pH 8.0.

The prepared 16HBE14o- cell lysate (sample volume used in the previous ^31^P NMR spectroscopy step) was diluted with 1 mL of dialysis buffer and added to a Vivaspin 20 protein concentrator spin column (3 kDa MWCO) and concentrated down to ~1.0 mL by centrifugation (6 °C, 3220 RCF). Dialysis buffer (1mL) was added, and again the mixture was concentrated down to ~ 1.0 mL as described above. This step was repeated 3 times. The mixture was concentrated down to the starting sample volume and 1/50 (wt/wt) trypsin was added. Once the trypsin had dissolved, the solution was mixed using an Eppendorf Thermomixer (600 rpm) at 37 °C for 16 hours. The sample was divided in two and to one half of the sample sodium carbonate (5 mg), sodium bicarbonate (0.5 mg) and 10 % (v/v) D_2_O was added and analysed by ^31^P NMR spectroscopy.

A C-18 resin Sep-Pak cartridge (130 mg) was equilibrated with acetonitrile (2.6 mL) followed by aqueous 50 % (v/v) acetonitrile + 0.1 % (v/v) trifluoroacetic acid (1.5 mL) and finally aqueous 10 % (v/v) trifluoroacetic acid. Trypsinized proteins from 16HBE14o- cell lysate were made up to 1 mL using water. The sample was acidified to pH 3 using aqueous 10 % (v/v) trifluoroacetic acid and passed through the equilibrated Sep-Pak cartridge. The flow through was collected and passed through the cartridge again, this step was repeated once more. The resin was washed with ice cold ultrapure water (2.6 mL) and the peptides were eluted from the resin using aqueous 50 % (wt/wt) acetonitrile (600 μL). The solvent was removed by speed vac at room temperature and the remaining residue was solubilized in 0.1 M carbonate/bicarbonate, pH 10.8 (same as starting volume) and analyzed by ^31^P NMR spectroscopy after the addition of 10 % (v/v) D_2_O.

### Trypsinization of proteins from the 16HBE14o- cell lysate and desalting under milder conditions

The method followed Hardman *et al*.^12^ with minor changes.

Lysis buffer: 50 mM ammonium bicarbonate, 8 M urea, complete protease inhibitor cocktail, pH 8.0.

The medium was removed from five cell culture dishes (57 cm^2^) containing adhered 16HBE14o- cells between 70-90 % confluent. The medium was discarded, and the cells were washed with PBS (10 mL). The petri dishes were cooled on ice and care was taken to remove as much PBS as possible. Ice cold lysis buffer (150 μL) was added to each petri dish. Each petri dish was carefully shaken to evenly spread the lysis buffer. The cells were scraped into the lysis buffer and the lysate from each dish was transferred to another vial on ice. The lysis procedure was repeated once more for a total of 10 cell culture dishes. The combined cell lysate was sonicated on ice (35 % amplitude 10 × 5 sec bursts, 59 sec rest between each burst). The lysate was treated with dithiothreitol (7 mM) and shaken (600 rpm) at 30° C for 20 minutes. After allowing the mixture to cool to room temperature, iodoacetamide (14 mM) was added and the mixture was left in the dark at room temperature. Dithiothreitol (3 mM) was added and the mixture was diluted to 2 M urea using 50 mM ammonium bicarbonate, pH 8.0. 2 % (wt/wt) trypsin was added and the mixture was shaken (600 rpm) at 30 ° C for 16 hours. A C-18 resin Sep-Pak cartridge (130 mg) was conditioned sequentially with methanol (2.6 mL), 50 % (v/v) acetonitrile (2.6 mL) and water (2.6 mL). The remaining sample (~500 μL) was made up to 2 mL using water and half of the sample (1 mL) was passed through the cartridge. The flow through was collected and passed through the cartridge again; this step was repeated once more. The resin was washed with water (2.6 mL), and then extracted with 50 % (v/v) acetonitrile (0.6 mL). The other half of the sample (1 mL) was desalted in the same way using a fresh cartridge. Both desalted samples where combined and solvent was removed under reduced pressure. The remaining residue was solubilized in 0.1 M sodium carbonate/hydrogen carbonate, pH 10.8 (500 μL) and analysed by ^31^P NMR spectroscopy after the addition of 10 % (v/v) D_2_O.

### DNA extraction and precipitation of proteins from the 16HBE14o- cell lysate

The method followed Antonioli *et al*. with slight modifications.^38^

Lysis buffers: 30 mm tris(hydroxymethyl)aminomethane-hydrochloride, 50 mm sodium chloride, complete protease inhibitor, pH 9.0.

Dialysis buffer: 8 M urea, 0.1 M sodium carbonate/bicarbonate, pH 10.8. DNA extraction solution: phenol/chloroform/isoamyl alcohol (25:24:1).

Lysed 16HBE14o- cells (700 μL, 18.1 mg of protein) were sonicated on ice (45% amplitude 10 × 5 sec bursts, 59 sec rest between each burst). The pH was raised to pH 10 using 2 M NaOH. The lysate was centrifuged to pellet any debris. The supernatant was removed and DNA extraction solution (1.4 mL) was added. The mixture was vortexed and left on ice for 5 minutes and then centrifuged (4 °C, 17000 RCF, 10 min). The upper aqueous and the bottom organic phase were carefully removed. Acetone (3ml) was added to the remaining precipitate. The precipitate was centrifuged, and the supernatant was removed. The precipitate was allowed to air dry and then solubilized in in dialysis buffer (1.5 ml). The mixture was added to a Vivaspin 20 protein concentrator spin column (3 kDa MWCO) and concentrated down to ~ 1.0 mL by centrifugation (6 °C, 3220 RCF, ~2.5 hr). 1 mL dialysis buffer was added, and the mixture was concentrated down as above to ~ 1.0 mL. This step was repeated 7 times. The sample was concentrated down to ~ 0.5 mL and 10 % (v/v) D_2_O was added before analysis by ^31^P NMR spectroscopy.

### Precipitation of proteins from the 16HBE14o- cell lysate

The method followed Potel *et al*.,^13^ with minor modification.

Lysed cells were prepared as described in “Preparation of proteins from the 16HBE14o- and Jurkat cell lysate for ^31^P NMR spectroscopy**”.** Total lysate volume after sonication: 6.5 mL. Methanol (13 mL), chloroform (3.3 mL), and water (10 mL) were sequentially added to the lysate and followed by vigorous vortexing after the addition of each solution. The mixture was centrifuged (6 °C, 3220 RCF, 10 min) and the upper layer was discarded. Methanol (10 mL) was added and the mixture was vortexed and then centrifuged (6 °C, 3220 RCF, 10 min). The supernatant was discarded. The precipitate was dissolved in 0.1 M sodium carbonate/bicarbonate, pH 10.8 and 10 % (v/v) D_2_O was added before ^31^P NMR spectroscopy.

### Chemical phosphorylation

Dialysis buffer: 8 M urea, 0.1 M sodium carbonate/bicarbonate, pH 10.8.

The proteins from the 16HBE14o- cells prepared from the “DNA extraction” procedure (500 μL) were treated with potassium phosphoramidate (68 mg, 0.5 mmol). The mixture was turned end over end at room temperature for 15 minutes. Dialysis buffer (400 μL) and water (200 μL) was added and the mixing was continued at room temperature for a further 1 hour. The mixture was made up to 3.5 mL with dialysis buffer. The mixture was added to a Vivaspin 20 protein concentrator spin column (3 kDa MWCO) and concentrated down to ~ 1.5 mL by centrifugation (6 °C, 3220 RCF, ~2.5 hr). 1 mL dialysis buffer was added, and the mixture was concentrated down as above to ~ 1.5 mL. This step was repeated 7 times. The sample finally was concentrated down to ~ 0.5 mL and 10 % (v/v) D_2_O was added before analysis by ^31^P NMR spectroscopy.

### Acid treatment of proteins from the 16HBE14o- cell lysate

The pH of proteins from the 16HBE14o- cell lysate in 91 mM sodium carbonate/bicarbonate, 6.4 M urea, 10 % (v/v) D_2_O was reduced to ~pH 4 using glacial acetic acid and it was then heated at 90 °C for 45 min. A precipitate was observed. Sodium carbonate (10 mg) and sodium hydrogen carbonate (1 mg) was added. 10 % (v/v) D_2_O was added to the sample before analysis by ^31^P NMR spectroscopy.

## Acknowledgements

We thank Dr C.C. Robertson for running ^31^P NMR samples, and EPSRC for support (DTA studentship to M.V.M.)

## References

1. Khoury GA, Baliban RC, Floudas CA. Proteome-wide post-translational modification statistics: Frequency analysis and curation of the swiss-prot database. Sci Rep. 2011;1:90–20. doi: 10.1038/srep00090 PMID: 22034591

2. Attwood PV, Piggott MJ, Zu XL, Besant PG. Focus on phosphohistidine. Amino Acids. 2007;32(1):145–56. doi: 10.1007/s00726-006-0443-6 PMID: 17103118

3. Besant PG, Attwood PV, Piggott MJ. Focus on phosphoarginine and phospholysine. Curr Protein Peptide Sci. 2009;10(6):536–50. doi: 10.2174/138920309789630598 PMID: 19751195

4. Attwood PV, Besant PG, Piggott MJ. Focus on phosphoaspartate and phosphoglutamate. Amino Acids. 2011;40(4):1035–51. doi: 10.1007/s00726-010-0738-5 PMID: 20859643

5. Piggott MJ, Attwood PV. Focus on o-phosphohydroxylysine, o-phosphohydroxyproline, n1-phosphotryptophan and s-phosphocysteine. Amino Acids. 2017;49(8):1309–23. doi: 10.1007/s00726-017-2446-x PMID: 28578504

6. Lad C, Williams NH, Wolfenden R. The rate of hydrolysis of phosphomonoester dianions and the exceptional catalytic proficiencies of protein and inositol phosphatases. Proc Natl Acad Sci USA. 2003;100(10):5607–10. doi: 10.1073/pnas.0631607100 PMID: 12721374

7. Hunter T. Why nature chose phosphate to modify proteins. Philos Trans R Soc Lond, Ser B: Biol Sci. 2012;367(1602):2513–6. doi: 10.1098/rstb.2012.0013 PMID: 22889903

8. Cohen P. The role of protein phosphorylation in human health and disease Eur J Biochem. 2001;268(19):5001–10. doi: 10.1046/j.0014-2956.2001.02473.x PMID: 11589691

9. Wadhams GH, Armitage JP. Making sense of it all: Bacterial chemotaxis. Nat Rev Mol Cell Biol. 2004;5(12):1024–37. doi: 10.1038/nrm1524 PMID: 15573139

10. Sickmann A, Meyer HE. Phosphoamino acid analysis. Proteomics. 2001;1(2):200–6. doi: 10.1002/1615-9861(200102)1:2<200::aid-prot200>3.3.co;2-m PMID: 11680867

11. Hornbeck PV, Zhang B, Murray B, Kornhauser JM, Latham V, Skrzypek E. Phosphositeplus, 2014: Mutations, ptms and recalibrations. Nucleic Acids Res. 2014;43(D1):D512–D20. doi: 10.1093/nar/gku1267 PMID: 25514926

12. Hardman G, Perkins S, Brownridge PJ, Clarke CJ, Byrne DP, Campbell AE, et al. Strong anion exchange-mediated phosphoproteomics reveals extensive human non-canonical phosphorylation. EMBO J. 2019;38:e100847–e. doi: 10.15252/embj.2018100847 PMID: 31433507

13. Potel CM, Lin M-H, Heck AJR, Lemeer S. Widespread bacterial protein histidine phosphorylation revealed by mass spectrometry-based proteomics. Nat Methods. 2018;15:187–90. doi: 10.1038/nmeth.4580 PMID: 29377012

14. Makwana MV, Muimo R, Jackson RFW. Advances in development of new tools for the study of phosphohistidine. Lab Invest. 2017;98:291. doi: 10.1038/labinvest.2017.126 PMID: 29200202

15. Klumpp S, Krieglstein J. Phosphorylation and dephosphorylation of histidine residues in proteins. Eur J Biochem. 2002;269(4):1067–71. doi: 10.1046/j.1432-1033.2002.02755.x PMID: 11856347

16. Adam K, Hunter T. Histidine kinases and the missing phosphoproteome from prokaryotes to eukaryotes. Lab Invest. 2017;98:233. doi: 10.1038/labinvest.2017.118 PMID: 29058706

17. Gonzalez-Sanchez M-B, Lanucara F, Hardman GE, Eyers CE. Gas-phase intermolecular phosphate transfer within a phosphohistidine phosphopeptide dimer. Int J Mass spectrom. 2014;367:28–34. doi: 10.1016/j.ijms.2014.04.015 PMID: 25844054

18. Cui L, Reid GE. Examining factors that influence erroneous phosphorylation site localization via competing fragmentation and rearrangement reactions during ion trap cid- ms/ms and -ms3. Proteomics. 2013;13(6):964–73. doi: 10.1002/pmic.201200384 PMID: 23335301

19. Rozman M. Modelling of the gas-phase phosphate group loss and rearrangement in phosphorylated peptides. J Mass Spectrom. 2011;46(9):949–55. doi: 10.1002/jms.1974 PMID: 21915960

20. Schmidt A, Ammerer G, Mechtler K. Studying the fragmentation behavior of peptides with arginine phosphorylation and its influence on phospho-site localization. Proteomics. 2013;13(6):945–54. doi: 10.1002/pmic.201200240 PMID: 23172725

21. Chen EI, Cociorva D, Norris JL, Yates JR, III. Optimization of mass spectrometry-compatible surfactants for shotgun proteomics. J Proteome Res. 2007;6(7):2529–38. doi: 10.1021/pr060682a PMID: 17530876

22. Gonzalez-Sanchez M-B, Lanucara F, Helm M, Eyers Claire E. Attempting to rewrite history: Challenges with the analysis of histidine-phosphorylated peptides. Biochem Soc Trans. 2013;41(4):1089–95. doi: 10.1042/bst20130072

23. Besant PG, Byrne L, Thomas G, Attwood PV. A chromatographic method for the preparative separation of phosphohistidines. Anal Biochem. 1998;258(2):372–5. doi: 10.1006/abio.1998.2595 PMID: 9570854

24. Himmel S, Wolff S, Becker S, Lee D, Griesinger C. Detection and identification of protein-phosphorylation sites in histidines through hnp correlation patterns. Angew Chem Int Ed. 2010;49(47):8971–4. doi: 10.1002/anie.201003965 PMID: 20939030

25. Hohenester UM, Ludwig K, Koenig S. Chemical phosphorylation of histidine residues in proteins using potassium phosphoramidate - a tool for the analysis of acid-labile phosphorylation. Curr Drug Del. 2013;10(1):58–63. doi: 10.2174/1567201811310010010 PMID: 22998046

26. Hultquist DE. Preparation and characterization of phosphorylated derivatives of histidine. Biochim Biophys Acta. 1968;153(2):329–40. doi: 10.1016/0005-2728(68)90078-9 PMID: 5642389

27. Kowalewska K, Stefanowicz P, Ruman T, Frączyk T, Rode W, Szewczuk Z. Electron capture dissociation mass spectrometric analysis of lysine-phosphorylated peptides. Biosci Rep. 2010;30(6):433–43. doi: 10.1042/bsr20090167 PMID: 20144148

28. Lecroisey A, Lascu I, Bominaar A, Veron M, Delepierre M. Phosphorylation mechanism of nucleoside diphosphate kinase - p-31-nuclear magnetic-resonance studies. Biochemistry. 1995;34(38):12445–50. doi: 10.1021/bi00038a043 PMID: 7547990

29. Muimo R, Banner SJ, Marshall LJ, Mehta A. Nucleoside diphosphate kinase and cl−-sensitive protein phosphorylation in apical membranes from ovine airway epithelium. Am J Respir Cell Mol Biol. 1998;18(2):270–8. doi: 10.1165/ajrcmb.18.2.2850 PMID: 9476915

30. Muimo R, Hornickova Z, Riemen CE, Gerke V, Matthews H, Mehta A. Histidine phosphorylation of annexin i in airway epithelia. J Biol Chem. 2000;275(47):36632–6. doi: 10.1074/jbc.M000829200 PMID: 10956639

31. Duclos B, Marcandier S, Cozzone AJ. Chemical-properties and separation of phosphoamino acids by thin-layer chromatography and or electrophoresis. Methods Enzymol. 1991;201:10–21. doi: 10.1016/0076-6879(91)01004-l PMID: 1943759

32. Fuhs SR, Meisenhelder J, Aslanian A, Ma L, Zagorska A, Stankova M, et al. Monoclonal 1- and 3-phosphohistidine antibodies: New tools to study histidine phosphorylation. Cell. 2015;162(1):198–210. doi: 10.1016/j.cell.2015.05.046 PMID: MEDLINE:26140597

33. Hoffmann R, Reichert I, Wachs WO, Zeppezauer M, Kalbitzer HR. ^1^H and ^31^P NMR spectroscopy of phosphorylated model peptides. Int J Pept Protein Res. 1994;44(3):193–8. doi: 10.1111/j.1399-3011.1994.tb00160.x PMID: 7529751

34. Robitaille P-ML, Robitaille PA, Gordon Brown G, Brown GG. An analysis of the ph-dependent chemical-shift behavior of phosphorus-containing metabolites. J Magn Reson. 1991;92(1):73–84. doi: 10.1016/0022-2364(91)90248-R

35. Cade-Menun BJ, Carter MR, James DC, Liu CW. Phosphorus forms and chemistry in the soil profile under long-term conservation tillage: A phosphorus-31 nuclear magnetic resonance study. J Environ Qual. 2010;39(5):1647–56. doi: 10.2134/jeq2009.0491 PMID: 21043270

36. Lilley M, Mambwe B, Thompson MJ, Jackson RFW, Muimo R. 4-phosphopyrazol-2-yl alanine: A non-hydrolysable analogue of phosphohistidine. Chem Commun. 2015;51(34):7305–8. doi: 10.1039/C5CC01811K PMID: 25820536

37. Martensen TM. Phosphotyrosine in proteins - stability and quantification. J Biol Chem. 1982;257(16):9648–52. PMID: 6179934

38. Antonioli P, Bachi A, Fasoli E, Righetti PG. Efficient removal of DNA from proteomic samples prior to two-dimensional map analysis. J Chromatogr. 2009;1216(17):3606–12. doi: 10.1016/j.chroma.2008.11.053 PMID: 19081104

39. Matthews HR. Protein-kinases and phosphatases that act on histidine, lysine, or arginine residues in eukaryotic proteins - a possible regulator of the mitogen-activated protein-kinase cascade. Pharmacol Ther. 1995;67(3):323–50. doi: 10.1016/0163-7258(95)00020-8 PMID: 8577821

40. Glonek T, Henderson TO, Hilderbrand RL, Myers TC. Biological phosphonates: Determination by phosphorus-31 nuclear magnetic resonance. Science. 1970;169(3941):192. doi: 10.1126/science.169.3941.192

41. Oslund RC, Kee J-M, Couvillon AD, Bhatia VN, Perlman DH, Muir TW. A phosphohistidine proteomics strategy based on elucidation of a unique gas-phase phosphopeptide fragmentation mechanism. J Am Chem Soc. 2014;136(37):12899–911. doi: 10.1021/ja507614f PMID: 25156620

42. Kiianitsa K, Maizels N. A rapid and sensitive assay for DNA-protein covalent complexes in living cells. Nucleic Acids Res. 2013;41(9):e104–e. Epub 03/21. doi: 10.1093/nar/gkt171 PMID: 23519618

43. Marmelstein AM, Yates LM, Conway JH, Fiedler D. Chemical pyrophosphorylation of functionally diverse peptides. J Am Chem Soc. 2014;136(1):108–11. doi: 10.1021/ja411737c PMID: 24350643

44. Zhou H, Di Palma S, Preisinger C, Peng M, Polat AN, Heck AJR, et al. Toward a comprehensive characterization of a human cancer cell phosphoproteome. J Proteome Res. 2013;12(1):260–71. doi: 10.1021/pr300630k

45. Hartsough MT, Morrison DK, Salerno M, Palmieri D, Ouatas T, Mair M, et al. Nm23-h1 metastasis suppressor phosphorylation of kinase suppressor of ras via a histidine protein kinase pathway. J Biol Chem. 2002;277(35):32389–99. doi: 10.1074/jbc.M203115200 PMID: 12105213

46. Walsh CT, Tu BP, Tang Y. Eight kinetically stable but thermodynamically activated molecules that power cell metabolism. Chem Rev. 2018;118(4):1460–94. doi: 10.1021/acs.chemrev.7b00510 PMID: 29272116

47. Chen C-W, Wang H-L, Huang C-W, Huang C-Y, Lim WK, Tu I-C, et al. Two separate functions of nme3 critical for cell survival underlie a neurodegenerative disorder. Proc Natl Acad Sci. 2019;116(2):566–74. doi: 10.1073/pnas.1818629116 PMID: 30587587

48. Postel EH, Zou X, Notterman DA, La Perle KMD. Double knockout nme1/nme2 mouse model suggests a critical role for ndp kinases in erythroid development. Mol Cell Biochem. 2009;329(1):45–50. doi: 10.1007/s11010-009-0110-9 PMID: 19381783

49. Kee JM, Oslund RC, Perlman DH, Muir TW. A pan-specific antibody for direct detection of protein histidine phosphorylation. Nat Chem Biol. 2013;9(7):416–21. Epub 2013/05/28. doi: 10.1038/nchembio.1259 PMID: 23708076

